# Sodium chloride in the tumor microenvironment enhances T-cell metabolic fitness and cytotoxicity

**DOI:** 10.1101/2023.09.14.557686

**Authors:** Dominik Soll, Mahima Arunkumar, Maha Alissa-Alkhalaf, Shan Sun, Trang Nguyen, Chang-Feng Chu, Veronika Lutz, Sascha Schäuble, Ignacio Garcia-Ribelles, Michael Mueller, Bernhard Michalke, Gianni Panagiotou, Philipp Schatzlmaier, Hannes Stockinger, Wolfgang W. Schamel, Magdalena Huber, Christina E. Zielinski

**Affiliations:** Technical University of Munich, 81675 Munich, Germany; Department of Infection Immunology, Leibniz Institute for Natural Product Research and Infection Biology, Hans Knöll Institute, 07745 Jena, Germany; Institute of Microbiology, Faculty of Biological Sciences, Friedrich Schiller University, 07745 Jena, Germany; Institute of Biology III, Faculty of Biology and Signalling Research Centres BIOSS and CIBSS, University of Freiburg, Freiburg, Germany; Center of Chronic Immunodeficiency (CCI), University Clinics and Medical Faculty, University, Freiburg, Germany; Spemann Graduate School of Biology and Medicine (SGBM), University of Freiburg, Germany; Institute of Systems Immunology, Philipps-University Marburg, 35043 Marburg, Germany; Department of Microbiome Dynamics, Leibniz Institute for Natural Product Research and Infection Biology, Hans Knöll Institute, 07745 Jena, Germany; Research Unit Analytical BioGeoChemistry, Helmholtz Center Munich - German Research Center for Environmental Health (GmbH), 85764 Neuherberg, Germany; Medical University of Vienna, Center for Pathophysiology, Infectiology and Immunology, Institute for Hygiene and Applied Immunology, 1090 Vienna, Austria

**Keywords:** human T cells, cytotoxicity, tumor microenvironment, T-cell metabolism

## Abstract

Adoptive T-cell therapy has become a powerful weapon for cancer treatment. The efficacy of antitumor immunity is associated with the metabolic state of cytotoxic T cells, which is highly sensitive to the tumor microenvironment. It is therefore of considerable interest to bypass immunosuppressive signals in the tumor microenvironment and to identify factors that augment cytotoxic effector functions and ultimately tumor killing. Whether ionic signals serve as aberrant immune signals and influence the adaptive human antitumor immune response is still largely unexplored. We therefore investigated the effect of sodium on the phenotype, function and metabolic regulation of human CD8^+^ T cells using transcriptomic, metabolomic, high-dimensional flow cytometric and functional assays. We demonstrate a significant enrichment of sodium in solid tumors from patients with breast cancer, which leaves a transcriptomic imprint on intratumoral immune cells. Sodium chloride (NaCl) enhanced the activation state and effector functions of human CD8^+^ memory T cells. These functional alterations were associated with enhanced metabolic fitness, particularly increases in glycolysis, oxidative phosphorylation and overall nutrient uptake. These NaCl-induced effects translated into increased tumor cell killing *in vitro* and in a tumor mouse model *in vivo.* We therefore propose NaCl as a positive regulator of acute antitumor immunity that could be harnessed for *ex vivo* conditioning of adoptively transferred T cells, such as CAR T-cells.

## Introduction

CD8^+^ T cells with reactivity against tumors can mediate potent cytotoxic effector responses, which rely on target specificity, potent effector functions and long-term maintenance. The tumor microenvironment is a critical participant in the neoplastic process by providing, on the one hand side, inflammatory cues that promote neoplastic transformation and, on the other side, immunosuppressive factors that paralyze the antitumor immune response. This poses a challenge for adoptive T cell therapies, whose clinical efficacy is limited by low infiltration of transferred T cells, their impaired T-cell maintenance and suppressed effector functions (DePeaux and Delgoffe, 2021).

T-cell differentiation, activation and function have been shown to correlate with distinct metabolic states as well as with the consumption of specific extracellular metabolites (Siska and Rathmell, 2015). Metabolic rewiring is required to support the increased biosynthetic demands of the effector functions and clonal expansion of CD8^+^ T-cells. Effector T cells and tumor cells display shared metabolic features, such as using aerobic glycolysis to produce energy (the Warburg effect) and enhanced dependence on glutamine (Pearce et al., 2013). Accordingly, experimental reduction in glucose availability leads to decreased production of the cytotoxic T-cell markers granzyme A, granzyme B and perforin by CD8^+^ T cells (Cham et al., 2008), highlighting glycolysis as a key component of cytotoxic T cell activation. Proteomic analysis demonstrated that cytotoxic T cells retained abundant amounts of the protein machinery components required for oxidative phosphorylation, suggesting that these cells have flexibility in meeting their metabolic demands (Hukelmann et al., 2016). Resting CD8^+^ memory cells possess a large spare respiratory capacity, defined as the difference between basal and maximal respiration, and increased mitochondrial mass, compared to their naïve circulating counterparts. This capacity licenses resting CD8^+^ memory cells for more effective expansion, cytokine production and thus recall responses upon antigen re-encounter (Geltink et al., 2018).

While cytokines and metabolites have been the main targets for achieving immunomodulation in the tumor microenvironment, several lines of investigation have recently suggested a role for extracellular ions in the modulation of T-cell effector responses. Potassium ions (K^+^) were found to be enriched in the necrotic tumor microenvironment and to suppress T-cell receptor (TCR)-driven T-cell effector programs, while concomitantly promoting the stemness and thus self-renewal, expansion and multipotency of these cells (Eil et al., 2016; Vodnala et al., 2019). This prompted the suggestion that K^+^ ions assume the role of a tumor-induced ionic checkpoint acting upon T-cell effector functions. Elevated sodium ion (Na^+^) concentration was demonstrated to strongly promote the differentiation of Th17 cells under polarizing conditions and to promote the upregulation of self-regulatory cytokines in these cells upon restimulation (Matthias et al., 2020; Wu et al., 2013) (Kleinewietfeld et al., 2013). Recently, Na^+^ ions were also shown to abrogate the immunosuppressive function of Treg cells through perturbation of mitochondrial respiration (Corte-Real et al., 2023; Hernandez et al., 2015). Mitochondrial Na^+^ influx via the Na^+^/Ca^2+^ exchanger (NCLX) blocked complex II/III of the electron transport chain, leading to decreased oxidative phosphorylation and thus decreased ATP production (Corte-Real et al., 2023). Additionally, Na^+^ ions promoted Th2 cell responses, presumably through mTORC2 signaling, which has implications for the pathogenesis of allergic diseases (Matthias et al., 2019b). The direct effect of Na^+^ ions on CD8^+^ T cells, and thus on antitumor cytotoxicity, however, remains unknown. In the present study, we found increased intratumoral Na^+^ concentrations with transcriptomic imprints on tumor-infiltrating immune cells. We demonstrate that increased extracellular Na^+^ concentrations enhance T cell metabolic fitness, T cell effector functions and tumor control.

## Results

### Sodium is increased in the tumor microenvironment and shapes the transcriptome of intratumoral CD8^+^ T cells

To assess the potential impact of ionic signals on the immune regulation of human CD8^+^ T cells in the tumor microenvironment, we first determined the concentrations of Na^+^ and K^+^ in the lesional and perilesional tissues of patients with breast cancer (**Figure 1A**). Fresh tissue biopsies were analyzed via inductively coupled plasma optical emission spectrometry (ICP–OES), which allows for the highly sensitive and specific detection of various elements in organic tissues (Fischereder et al., 2017). We found significantly higher concentrations of Na^+^ in the lesional breast cancer tissue than in patient-matched perilesional tissue (**Figure 1B**). Consistent with previous reports (Eil et al., 2016), we also observed a significant accumulation of K^+^ in the tumor tissue (**Figure 1C**).

**Figure 1.**
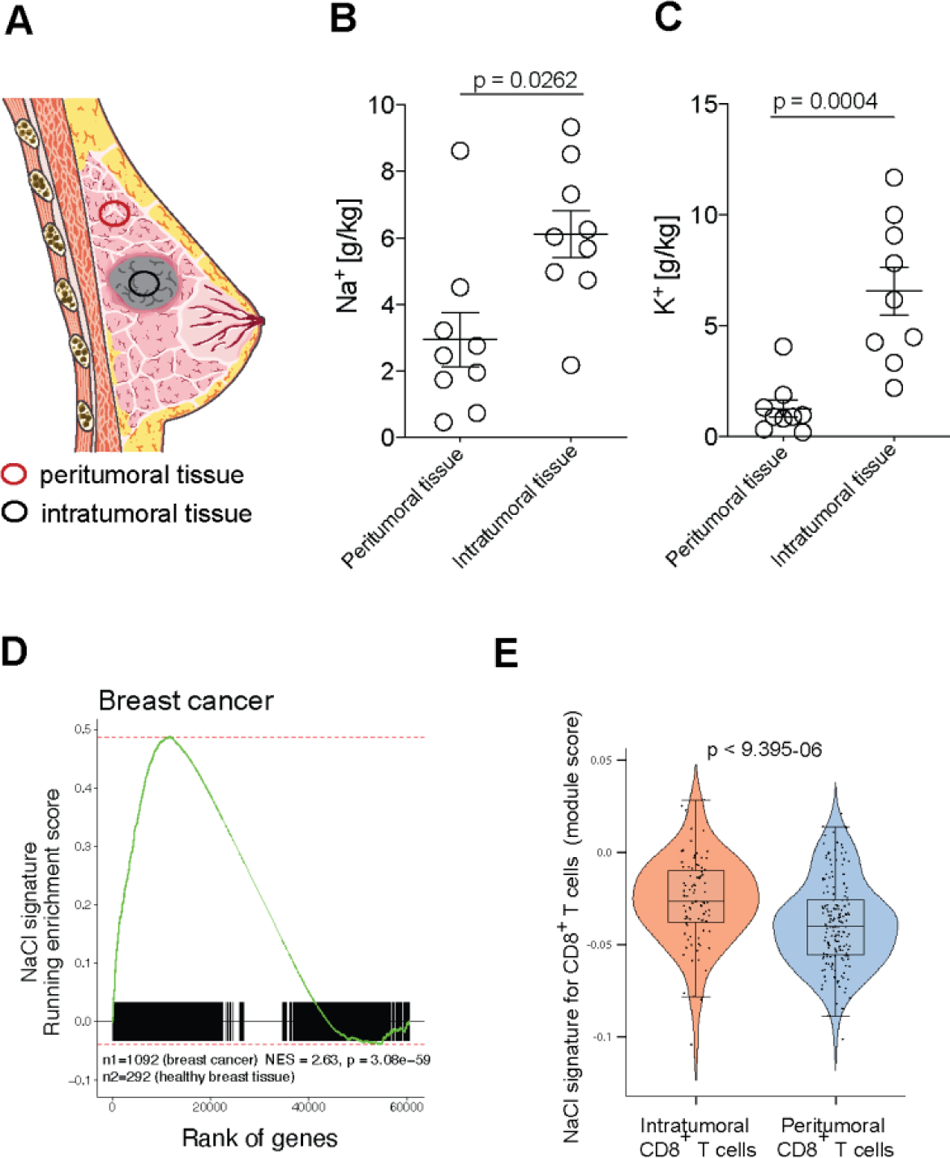
NaCl is highly enriched in solid tumors and leaves a transcriptomic imprint on infiltrating immune cells. **(A)** Schematic presentation of lesional and perilesional tissue biopsies from patients with breast cancer. **(B, C)** Quantification of Na^+^ (B) and K^+^ (C) concentrations with inductively coupled plasma optical emission spectrometry in intratumoral and peritumoral biopsies from patients with breast cancer. Paired Student’s t test. **(D)** GSEA of patients with breast cancer from The Cancer Genome Atlas (TCGA). Running enrichment score and sorted position of the 1956 significantly upregulated genes from the NaCl signature are shown. The NaCl signature was generated by transcriptomic comparison of CD8^+^ memory T cells cultured under high versus low NaCl conditions (top 60 significantly upregulated DEGs). NES: normalized enrichment score, p: significance of the enrichment (one-tailed test for positive enrichment), n_1/2_ = number of patients providing solid tumor samples or providing healthy breast tissue samples, respectively. **(E)** scRNA-seq of intra- and peritumoral immune cells from patients with breast cancer (n=3). The module score for the transcriptomic NaCl signature (obtained as described in D) was tested in intra- and peritumoral CD8^+^ T cells, which were identified by marker gene expression.

We next wished to explore whether the increased Na^+^ concentrations in the intratumoral microenvironment might impact antitumor immunity. We therefore investigated whether total tumor tissue, and specifically tumor-infiltrating human CD8^+^ T cells, displayed a transcriptomic scar after NaCl exposure. To this end, we first characterized the transcriptomic signature associated with NaCl exposure for CD8^+^ T cells by stimulating memory CD8^+^ T cells with CD3 and CD28 mAbs for 5 days under low (standard 140mM) versus high (+ 67.5 mM) extracellular NaCl conditions *in vitro*. Differentially expressed genes (DEGs), which were significantly upregulated following high-NaCl treatment, served as the gene set (NaCl signature) for further downstream analyses. 36 to 1278 patients per cancer type (14171 samples of 9554 patients in total) and 25 distinct tumor types from the publicly available Cancer Genome Atlas (TCGA) and Genotype-Tissue Expression (GTEx) projects were investigated to evaluate the enrichment of the NaCl signature in cancer tissues compared to normal tissue counterparts. Interestingly, we observed that breast cancer and all other solid tumor entities with the exception of tumors in only the thymus and paraganglia, displayed a significant enrichment of the NaCl signature relative to paired healthy tissues (23 out of 25, **Figure 1D, Figure S1)**.

Given that we found the identified transcriptomic NaCl signature to be enriched in whole tissue samples, which include all cells that comprise the tumor microenvironment, we next sought to determine whether this same signature would be evident in tumor-infiltrating CD8^+^ T cells (CD8^+^ TILs). To achieve this, we analyzed public single-cell RNA-seq (scRNA-seq) data from matched tumoral and peritumoral tissues from patients with breast cancer (Azizi et al., 2018) and determined the module score for the NaCl signature after filtering for tissue-infiltrating tumoral and peritumoral CD8^+^ T cells **(Figure S2A-C)**. In all three patients tested, we found a stronger enrichment of the NaCl signature in CD8^+^ TILs than in matched T cells from the perilesional breast tissue (**Figure 1E**).

Together, these data reveal that Na^+^ enrichment, a characteristic feature of the tumor microenvironment in patients with breast cancer and many other solid tumors, significantly shapes the transcriptome of tumor-infiltrating CD8^+^ T cells in humans.

### NaCl increases T-cell activation and TCR signaling in human CD8^+^ T cells

To gain mechanistic insight into the effects of NaCl on CD8^+^ T cells, we next explored the specific effect of NaCl on the bulk transcriptome of human memory CD8^+^ T cells *in vitro*. First, we used principal component analysis (PCA) to show that high NaCl conditions significantly changed the overall transcriptome of CD3- and CD28-stimulated human memory CD8^+^ T cells in three healthy donors (**Figure 2A**). A total of 1956 genes were significantly up- and 1926 genes were significantly down-regulated, respectively. Among these, we observed that *SGK1*, which encodes a NaCl-inducible kinase, was significantly upregulated upon stimulation under high NaCl conditions in CD8^+^ memory T cells (**Figure 2B**). These results are consistent with those previously observed in macrophages and T helper cells (Matthias et al., 2019b; Müller et al., 2019; Wu et al., 2013). Additionally, several genes encoding members of the solute carrier group of membrane transport proteins were among the top 50 significantly upregulated DEGs (*SLC5A3, SLC35F3, SLC12A8,* and *SLC29A1*) indicating enhanced uptake of various metabolites, such as sugar (*SLC5A3*) and amino acids (*SLC7A5*) and to glycolytic metabolic processing (*HK1, HK2*) upon exposure to high NaCl (**Figure 2B**). Notably, *BATF3*, which augments CD8^+^ T-cell metabolic fitness, viability and memory development, was among the top upregulated genes (Ataide et al., 2020) (**Figure 2B**). Additionally, *LTA,* which encodes lymphotoxin alpha, a cytotoxic effector protein with implications in tumor killing (Gray et al., 1984), was significantly upregulated by NaCl (**Figure 2B**). Likewise, *IRF4* expression, which is crucial for the sustained expansion and effector function of cytotoxic CD8^+^ T cells, was significantly increased under high NaCl conditions (Yao et al., 2013). Together, these data demonstrate NaCl-induced changes in the transcriptome of human CD8^+^ memory T cells, which point to an enhanced state of activation.

**Figure 2.**
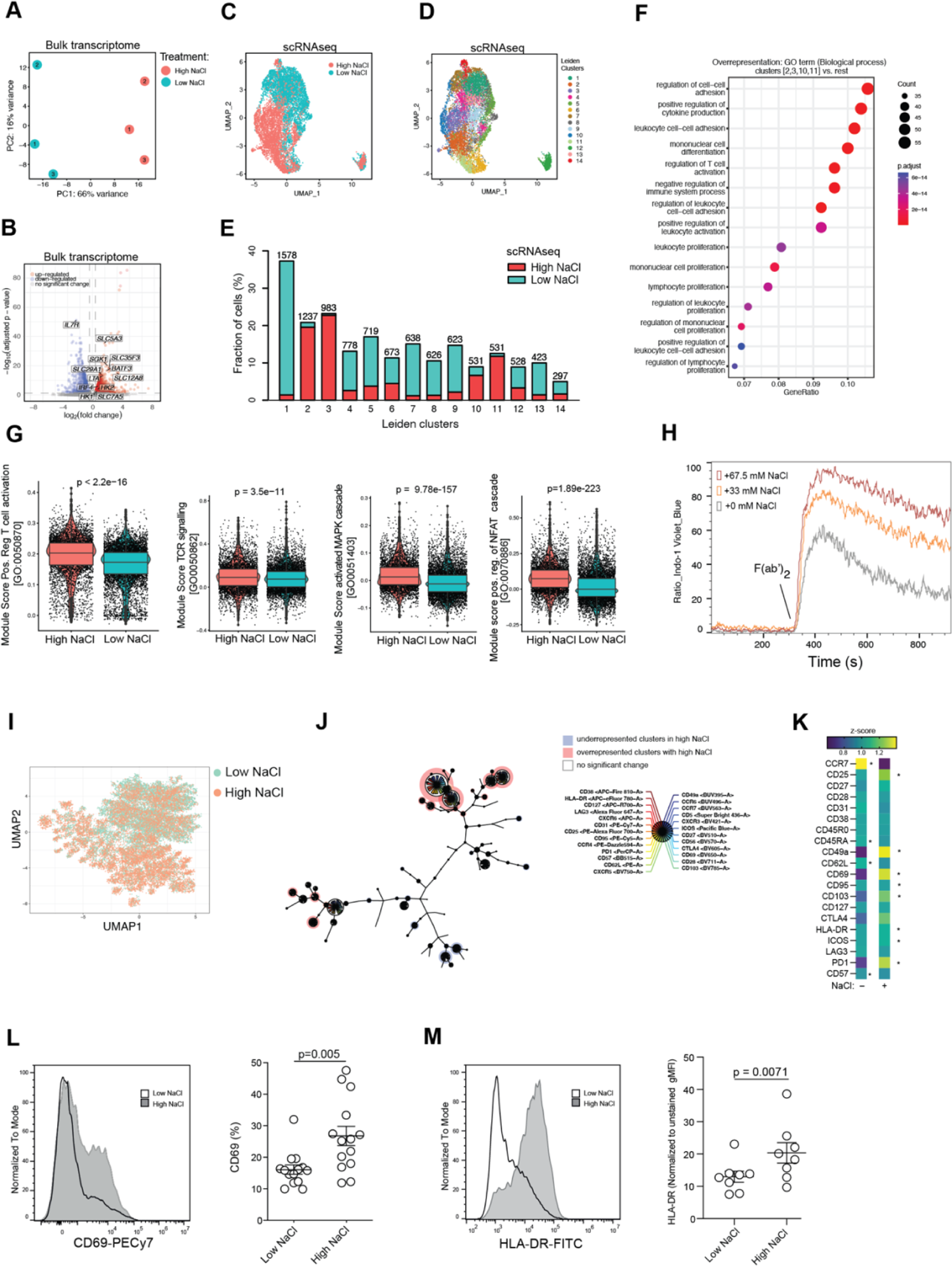
NaCl enhances the activation state of human CD8^+^ memory T cells. **(A)** PCA projection after mRNA-seq of bulk human CD8^+^ memory T cells stimulated with CD3 and CD28 mAb for 5 days under high versus low NaCl conditions. Numbers 1-3 represent individual blood donors. **(B)** mRNA-seq analysis of bulk human CD8^+^ memory T cells stimulated as in (A). Selected significant DEGs are highlighted in the Volcano plot visualization. **(C)** scRNA-seq of human memory CD8^+^ T cells stimulated with CD3 and CD28 mAb for 3 days under high versus low NaCl conditions. Shown is a UMAP representation visualizing the distribution of cells according to their respective treatment condition. **(D)** UMAP representation of CD8^+^ memory T cells stimulated as in (C) demonstrating Leiden clustering. **(E)** Distribution of CD8^+^ memory T cells from high versus low NaCl conditions within the Leiden clusters following scRNA-seq as in (C). **(F)** GSEA using the clusterProfiler enrichGO() function with all significantly upregulated DEGs from scRNA-seq comparing CD8^+^ memory T cells from high-salt clusters (2, 3, 10, and 11) versus the remaining low-salt Leiden clusters (D). The top 15 GO terms among all biological processes are shown. **(G)** Module scores for the indicated gene sets comparing CD8^+^ memory T cells from high versus low NaCl conditions after scRNA-seq. The Wilcoxon rank sum test was used. **(H)** Ca^2+^ flux measurement in real time by flow cytometry. Human CD8^+^ memory T cells were stimulated with CD3 and CD28 mAbs for 5 days in low and titrated high NaCl conditions (+33 mM, +67.5 mM), then washed before baseline Ca^2+^ flux measurements and measurements after TCR-crosslinking with an CD3 mAb and anti-mouse IgG (F(ab)’_2_. Representative data, n=6 blood donors. **(I)** Spectral flow cytometry with 33 protein surface markers of CD8^+^ memory T cells on day 5 after activation with CD3 and CD28 mAb in the indicated conditions. Dimensionality reduction visualization with UMAP highlighting the high versus low NaCl conditions according to the colour code. n=4 blood donors. **(J)** Minimum spanning tree of the 100 clusters and 22 metaclusters identified after FlowSOM analysis. The node size corresponds to the number of cells included and the size of each pie chart segment represents the median expression of each marker represented in rainbow colors for each cluster. Clusters overrepresented in the high salt condition are highlighted in red, and those underrepresented in high salt are colored in blue. n=4, significance was determined by the Wilcoxon rank sum test with a fold change threshold of 2.5. **(K)** Spectral flow cytometry as in (I). The differences in the percentages of cells positive for the shown markers were visualized by z-score. (n=4). *, p<0.05. Paired t test. * is located on the treatment side (high or low NaCl) that shows significant upregulation **(L, M)** Flow cytometry analysis of the indicated activation markers of human CD8^+^ memory T cells from independent donors after stimulation for 5 days with CD3 and CD28 mAbs under high versus low NaCl conditions. Paired Student’s t test.

Taking the heterogeneity of CD8^+^ memory T cells into account, we then performed a scRNA-seq analysis of CD45RA^-^CD8^+^ T cells that were stimulated for 3 days with CD3 and CD28 mAbs under high and low NaCl conditions to dissect differential effects of NaCl on distinct CD8^+^ T-cell types. Dimensionality reduction by Uniform Manifold Approximation and Projection (UMAP) supported our conclusion that memory CD8^+^ T cells treated under high versus low NaCl conditions clustered separately (**Figure 2C**). Leiden clustering was performed to reveal the heterogeneity within the memory CD8^+^ T cell population and its differential response to NaCl. It demonstrated 14 individual clusters (**Figure 2D**). We then we tested the overall viability of CD8^+^ memory T cells upon stimulation with a wide range of extracellular NaCl concentrations over a 5-day culture period. We found that it was neither altered by high as compared to low NaCl concentrations as determined by total cell numbers and cell cycles, nor did the frequency of dying or apoptotic cells change in response to high NaCl conditions **(Figure S3A-C)**. Interestingly, CD8^+^ T cells from high NaCl conditions dominated only 4 clusters numerically (cluster 2, 3, 10, 11), comprising the majority all CD8^+^ T cells (61%). In contrast, matched CD8^+^ T cells from low NaCl conditions, clustered much more distinctly and distributed over 10 individual clusters, instead (**Figure 2E**). Together, this indicates that NaCl exerted a dominant effect on the overall transcriptome of CD8^+^ T cells, thereby reducing its heterogeneity on the single-cell level within the CD8^+^ memory T cell population.

Unbiased overrepresentation analysis with differentially expressed genes (DEGs) from these high NaCl clusters (2, 3, 10, 11) demonstrated that activation, proliferation, differentiation and effector functions, which are known to be relevant for antitumor immunity, were among the top 15 enriched biological processes (**Figure 2F**). This finding was supported with a gene overrepresentation analysis by comparison of CD8^+^ T cells from either high or low NaCl conditions **(Figure S4)**. We found a significantly increased expression of the module score for T cell activation on the single-cell transcriptomic level, as well as for pathways representing proximal T cell receptor activation and activatory signalling downstream of the TCR such as the MAPK and NFAT pathways (**Figure 2G**).

To assess a potential functional effect of NaCl on proximal TCR signaling, we quantified TCR-stimulation-induced calcium influx (Ca^2+^ flux) (Pores-Fernando and Zweifach, 2009). To this end human CD45RA^-^ CD8^+^ T cells were expanded with CD3 and CD28 mAbs for 5 days under low and increasing NaCl concentrations (+ 0 mM, + 33 mM, + 67.5 mM) before Ca^2+^ flux was assessed by flow cytometry upon TCR restimulation using CD3 mAbs and a secondary F(ab)’_2_ fragment in the absence of additional NaCl (Pores-Fernando and Zweifach, 2009). Interestingly, we observed significantly elevated Ca^2+^ flux upon TCR restimulation if CD8^+^ T cells were preactivated in high NaCl microenvironments in a dose dependent manner (**Figure 2H**). These data demonstrate that a recent history of NaCl exposure can rewire TCR ligation in human CD8^+^ T cells to translate into increased downstream signaling.

Factors that amplify TCR signaling in response to a given tumor antigen invigorate T cells for improved effector functions and might reverse a blunted T-cell response. We therefore next aimed to determine that whether NaCl potentiates T cell activation on the protein level. We performed high-dimensional spectral flow cytometry for multiple T cell activation and differentiation associated protein surface markers (33 marker panel) for CD8^+^ memory T cells stimulated under high and low NaCl conditions. UMAP dimensionality reduction analysis demonstrated distinct clustering of CD8^+^ T cells exposed to high and low NaCl conditions (**Figure 2I**), similar to that we had observed on the transcriptomic level before. FlowSOM analysis highlighted the differential effect of NaCl within the human CD8^+^ memory T cell population. This analysis revealed significant overrepresentation of cells from the high NaCl conditions in 19 clusters (23.01% of all cells) and 19 significantly underrepresented clusters (22.10% of cells). 54.88% of all cells (62 clusters) remained resilient to the effects of NaCl on day 5 of stimulation (**Figure 2J**). Based on the expression of various early and late activation markers, the differences in protein marker expression between CD8^+^ T cells subjected to high and low NaCl conditions were partially attributable to significantly elevated activation states in response to high NaCl (**Figure 2K**). In particular, CD69, a classical marker of early leukocyte activation, was highly upregulated under high NaCl conditions (**Figure 2L**). This pattern also applied to HLA-DR, another marker of human T-cell activation and a proxy for recent cell division **and cytotoxic effector functions** (Tippalagama et al., 2021) (**Figure 2M**). Other activation-associated molecules such as ICOS (*ICOS*), CD103 (ITGAE) and PD-1 (PDCD1), were also significantly upregulated at the protein and transcriptomic levels in response to TCR activation under high NaCl as compared to low NaCl conditions (**Figure 2K**). Taken together, these data suggest that high NaCl conditions potentiate CD8^+^ T-cell activation at the transcriptomic, TCR-signaling and functional levels.

### NaCl potentiates cytotoxic T-cell effector functions

CD8^+^ T cells exert their effector functions through paracrine delivery of proapoptotic cytotoxic effector molecules, such as granzymes, via perforin pores. Given the increased NaCl concentrations in the tumor microenvironment and their impact on the overall CD8^+^ T-cell transcriptome *in vivo*, we investigated the effect of NaCl on specific CD8^+^ T-cell effector functions. Transcriptome-wide analysis of bulk memory CD8^+^ T cells that were stimulated with CD3 and CD28 mAb under high and low NaCl conditions, demonstrated a significant shift toward the effector cell identity paired with reduced stemness upon stimulation under high NaCl conditions (**Figure 3A**). To explore the impact of NaCl on the differentiation state of CD8^+^ T cells in more detail on the single-cell level, we performed a trajectory inference analysis following scRNA-seq using RNA velocity, which can predict the future state of individual cells by distinguishing between unspliced and spliced single-cell mRNAs (La Manno et al., 2018). RNA velocity demonstrated a clear of differentiation trajectory for CD8^+^ T cells from low to high salt conditions (**Figure 3B**). This finding was supported by higher expression of the effector function (Pauken et al., 2016) (**Figure 3C**) and cytokine activation modules (**Figure 3D**) in CD8^+^ T cells from high NaCl conditions than in CD8^+^ T cells from low NaCl conditions.

**Figure 3.**
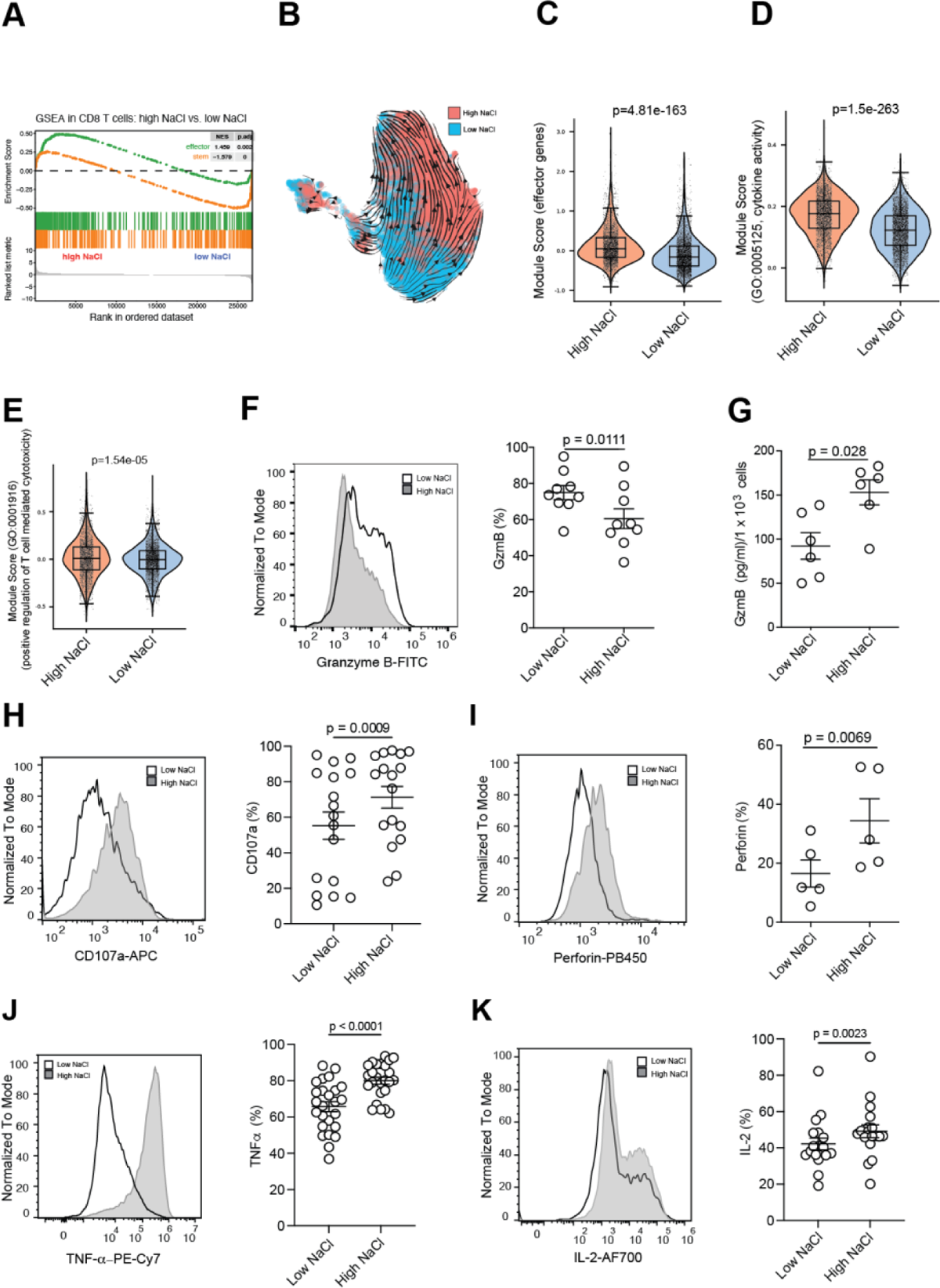
NaCl enhances CD8^+^ T-cell effector functions and cytotoxicity. **(A)** GSEA for effector and stemness associated genes was performed after a transcriptomic comparison of bulk human CD8^+^ memory T cells stimulated with CD3 and CD28 mAbs for 5 days under high versus low NaCl conditions. **(B)** RNA velocity analysis following scRNA-seq of human memory CD8^+^ T cells stimulated with CD3 and CD28 mAb for 3 days under high versus low NaCl conditions. Shown are the velocities in UMAP embedding. Cells are colour coded according to treatment condition as indicated (**C-E)** scRNAseq and analysis of the module scores of the indicated gene sets for human CD8^+^ memory T cells stimulated with CD3 and CD28 mAbs for 3 days under high and low NaCl conditions. Wilcoxon rank sum test. **(F)** Intracellular cytokine staining and flow cytometric analysis of human CD8^+^ memory T cells after stimulation for 5 days with CD3 and CD28 mAbs for 5 days under high and low NaCl conditions. Left, representative experiment. Right, cumulative data. Paired Student’s t test. **(G)** ELISA of cell culture supernatants from human CD8^+^ memory T cells stimulated for 5 days with CD3 and CD28 mAbs. **(H)** Flow cytometric analysis of human CD8^+^ memory T cells was performed after stimulation for 5 days with CD3 and CD28 mAbs under high versus low NaCl conditions. Left, representative experiment. Right, cumulative data. Paired Student’s t test. **(I-K)** Intracellular cytokine staining and flow cytometric analysis of human CD8^+^ memory T cells after stimulation for 5 days with CD3 and CD28 mAbs under high and low NaCl conditions and restimulation with PMA/ionomycin for 5 h. Left, representative experiment. Right, cumulative data. Paired Student’s t test.

Notably, CD8^+^ T cells from high NaCl conditions also displayed an increased module score for the GO term “positive regulation for T-cell mediated cytotoxicity” (**Figure 3E**). We then specifically analyzed the expression of granzyme B, one of the most abundant granzymes in cytoplasmic granules of human T cells. Intracellular granzyme B protein expression was significantly reduced upon CD8^+^ T-cell stimulation under high NaCl conditions on day 5 (**Figure 3F**). This finding reflected strong release of preformed granzyme B into the extracellular space, as demonstrated by ELISA of cell culture supernatants (**Figure 3G**) and by significant upregulation of the degranulation marker CD107a and perforin (**Figure 3H, I)**. Together, these results indicate that NaCl potentiates the cytotoxic effector functions of human CD8^+^ T cells.

To further explore the functionality of human CD8^+^ in high NaCl conditions, we next tested the effect of NaCl on CD8^+^ T-cell associated cytokines. In addition to its proinflammatory effects, TNF-α is known to induce cytotoxicity in select target cells (Ratner and Clark, 1993). TNF-α expression was significantly increased in CD8^+^ T cells under high compared to low NaCl conditions (**Figure 3J**). Likewise, we found increased production of IL-2 by CD8^+^ T cells in high NaCl conditions, consistent with the enhanced activation and effector differentiation of these cells in high NaCl conditions **(Figure K)**. Together, these results demonstrate that NaCl potentiates the cytotoxic effector functions of human CD8^+^ memory T cells.

### NaCl boosts CD8^+^ T-cell metabolic fitness

Given our findings that exposure to increased extracellular NaCl concentrations translated into increased CD8^+^ T-cell activation and cytotoxic effector functions, we considered the possibility of metabolic reprogramming of T cells by NaCl as a potential underlying mechanism. T cells are known to undergo a metabolic switch upon engagement of TCR signaling. This metabolic switch involves increases in nutrient uptake, aerobic glycolysis, glutaminolysis and lipid synthesis to support the execution of potent T-cell effector functions (Pearce et al., 2013). In line with this hypothesis, an unbiased bulk transcriptomic comparison of memory CD8^+^ T cells stimulated under high as compared to low NaCl conditions demonstrated overall increases in a multitude of cell metabolism associated processes among all upregulated KEGG pathways (**Figure 4A**). We then assessed the overall production of ATP, the principal energy currency of each cell (Pearce et al., 2013). As expected, in flow cytometric analysis, we observed increased intracellular ATP levels at the single-cell level in human memory CD8^+^ T cells subjected to polyclonal activation with CD3 and CD28 mAbs compared to naive CD8^+^ T cells (van der Windt et al., 2013). Interestingly, increased concentrations of extracellular NaCl resulted in a significant increase in intracellular ATP (**Figure 4B**).

**Figure 4.**
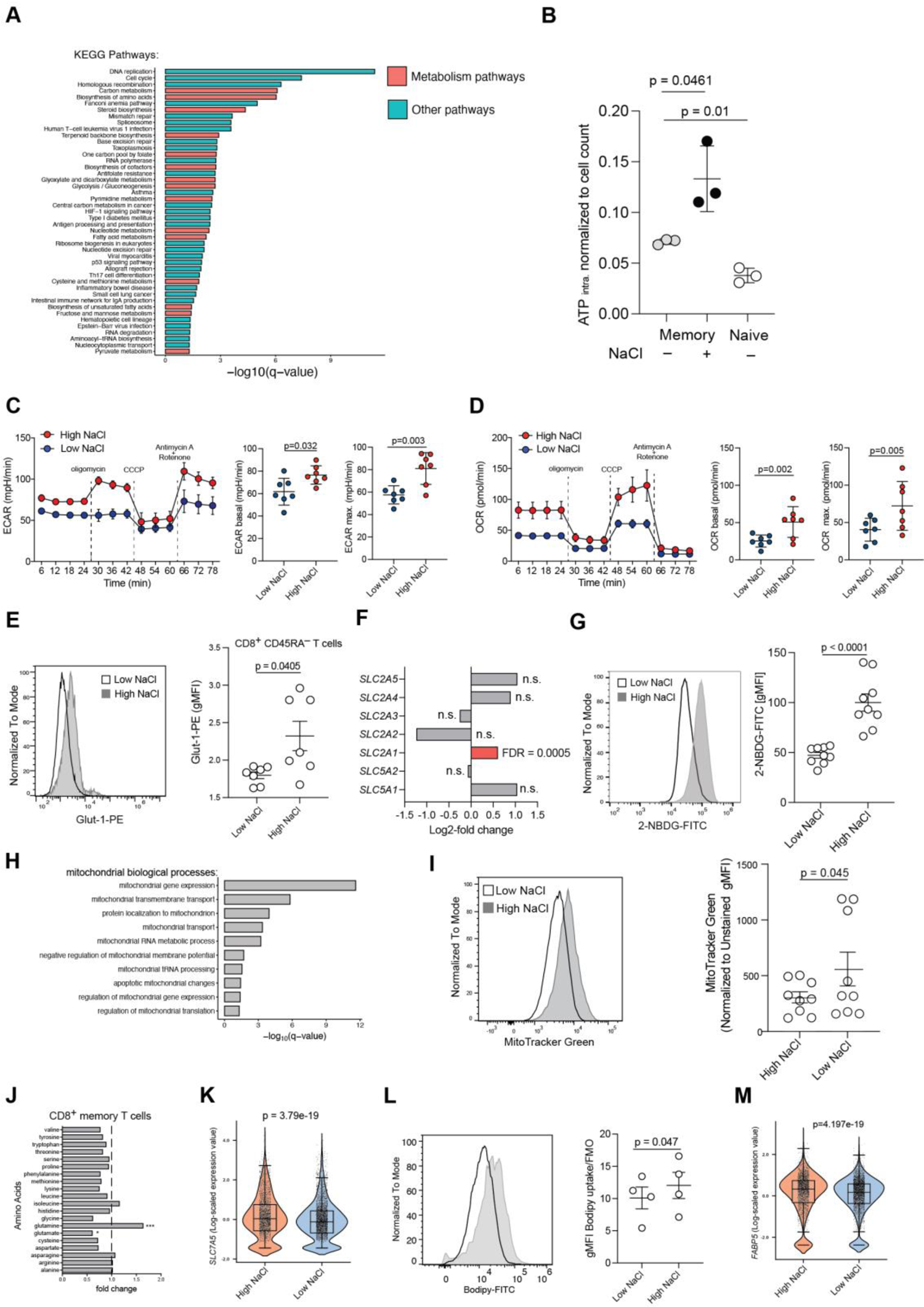
NaCl potentiates the metabolic fitness of CD8^+^ T cells. **(A)** Overrepresentation analysis for KEGG pathways (with false discovery adjusted q-value ≤ 0.05) using significantly upregulated DEGs (p value ≤ 0.05, log2(fold change) ≥ 0.5) from the bulk transcriptomic comparison of human CD8^+^ memory T cells stimulated for 5 days with CD3 and CD28 mAbs under high versus low NaCl conditions. All 45 upregulated KEGG pathways are displayed. KEGG pathways representing metabolic pathways are shown in red, and other pathways are shown in blue. **(A)** Luminometric assessment of ATP production in human naïve CD45RA^+^CD8^+^ and memory CD45RA^-^CD8^+^ cells, which were stimulated as described in (A) under high and low NaCl conditions **(C, D)** Real-time analysis of the ECAR (C) and OCR (D) in human CD8^+^ memory T cells under the indicated conditions. Dotted lines show the time point of addition of the indicated substances. Left, representative experiment with technical replicates. Right, cumulative quantification with individual healthy donors. Mean ± S.E.M. Paired Student’s t test **(E, G, I, L)** Flow cytometric analysis of human CD8^+^ memory T cells under the indicated conditions. Left, representative experiment. Right, cumulative quantification. Paired Student’s t test. **(F)** Expression of the indicated genes encoding glucose transporters in a transcriptomic comparison of bulk human CD8^+^ memory T cells stimulated for 5 days with CD3 and CD28 mAbs under high and low NaCl conditions. **(H)** Significantly enriched terms of GO-annotated mitochondrial biological processes following overrepresentation analysis of upregulated genes from the bulk transcriptomic comparison of human CD8^+^ memory T cells stimulated as in (A). q-values show p value adjustment for multiple test correction. Term redundancy was reduced using REVIGO (http://revigo.irb.hr, similarity parameter = 0.5). **(J)** Nontargeted metabolic profiling (metabolome analysis) of memory CD8^+^ T cells after 5 days of stimulation with CD3 and CD28 mAbs under high versus low NaCl conditions. n= 4 individual blood donors (matched samples). One-way ANOVA. **(K, M)** scRNA-seq analysis. Wilcoxon rank sum test.

Glycolysis and oxidative phosphorylation (OXPHOS) are the two main cooperative processes that supply ATP within cells (Pearce et al., 2013). We therefore monitored the extracellular acidification rate (ECAR) and the cellular oxygen consumption rate (OCR) in CD8^+^ T cells in real-time as measures of mitochondrial respiration and glycolysis, respectively, following 5 days of stimulation with CD3 and CD28 mAbs under high or low NaCl conditions. We found substantially increased ECAR (**Figure 4C**) and OCR (**Figure 4D**) values in CD8^+^ memory T cells that were preconditioned with high NaCl.

These findings prompted an in-depth dissection of the metabolic perturbations induced by NaCl. We first focused on glycolysis. We demonstrated a significant overrepresentation of the glycolysis pathway by performing a transcriptomic comparison of bulk CD8^+^ memory T cells cultured under high versus low NaCl conditions (**Figure 4A**). Furthermore, 18 out of 20 genes involved in the glycolysis pathway were significantly upregulated in the KEGG pathway analysis and GSEA **(Figure S5A, B)**. We then investigated the expression of the glucose transporter Glut-1 by flow cytometry (**Figure 4E**) (Macintyre et al., 2014). High NaCl conditions resulted in significantly increased Glut-1 expression on the cell surface suggesting improved glucose uptake (**Figure 4E**). This finding was also consistent with the transcriptional upregulation of *GLUT1* (*SLC2A1*) under high NaCl conditions (**Figure 4F).** The expression of other glucose transporter genes, however, remained unaffected by NaCl (**Figure 4F**). We then confirmed by flow cytometric analysis of CD8^+^ memory T cells, which were stimulated with CD3 and CD28 mAb under high and low NaCl conditions, that increased concentrations of extracellular NaCl could indeed enhance cellular glucose uptake by promoting the influx of 2-NBDG, a fluorescent derivative of glucose (Yamada et al., 2007) (**Figure 4G**).

Given increased ECAR rates, we next focused on mitochondrial metabolism of CD8^+^ T cells in response to high NaCl conditions. Na^+^ has previously been shown to act as a second messenger with effects on OXPHOS function and redox signaling via the NCLX Na^+^/Ca^2+^ exchanger (Hernansanz-Agustin et al., 2020). Transcriptomic enrichment analysis showed upregulation of multiple functional terms associated with mitochondrial metabolism in response to high NaCl concentrations (**Figure 4H**). Analysis of MitoTracker Green staining by flow cytometry validated that CD8^+^ T-cell stimulation in high NaCl microenvironments resulted in increased mitochondrial mass, which was consistent with our observation of enhanced ATP production (**Figure 4I**). Additionally, metabolomic analysis using liquid chromatography and mass spectrometry demonstrated that out of 20 protein building amino acids only glutamine was significantly enriched in human memory CD8^+^ T cells upon stimulation under high NaCl conditions (**Figure 4J**). Glutamine is known to increase in abundance upon TCR activation and to support mitochondrial oxidative metabolism, acting as an alternative fuel for fatty acid synthesis (Altman et al., 2016; Carr et al., 2010; Hensley et al., 2013). The levels of glutamate, which has previously been shown to inhibit cytotoxic T cell differentiation (Johnson et al., 2018), were, accordingly, significantly decreased under high NaCl conditions (**Figure 4J**). Consistent with this finding, scRNA-seq analysis revealed increased expression of the *SLC7A5* glutamine transporter under high conditions (**Figure 4K**).

We finally focused on lipid metabolism. To this end, we tested uptake of the fluorescently labelled lipid analogue Bodipy by flow cytometry as a surrogate readout for lipid uptake by human CD8^+^ T cells. Interestingly, high NaCl concentrations induced significantly enhanced uptake of Bodipy by memory CD8^+^ T cells (**Figure 4L**). This finding was consistent with augmented *FABP5* expression upon transcriptomic comparison of single-cells stimulated in high versus low NaCl conditions (**Figure 4M**). Cumulatively, these analyses demonstrated an augmented uptake of nutrients in high NaCl conditions and profound metabolic rewiring of human CD8^+^ memory cells, which matches their increased activation and effector functions.

### NaCl promotes antigen-specific tumor cell killing by human CD8^+^ T cells

The increases in NaCl-induced effector functions and their metabolic correlates prompted an investigation into the overall killing capacity of human CD8^+^ T cells in the context of elevated NaCl concentrations. We therefore generated antigen-specific CD8^+^ memory T cells via nucleofection of a MART-1-specific TCR. Cells that stained positive or negative for the MART-1 TCR after nucleofection, as assessed by detection of the concomitantly introduced murine TCR β-chain **(Figure S6A, B)**, were stimulated under high versus low NaCl conditions before coculture with adherent MART-1 expressing A375 melanoma cells at a 1:1 ratio. Stable maintenance of the newly introduced TCR in cells subjected to both extracellular NaCl concentrations was confirmed by flow cytometry **(Figure S6A, B)**. Killing of tumor cells was monitored in real time using xCELLigence technology. We observed a strong increase in specific melanoma target cell lysis by CD8^+^ T cells that were prestimulated for 5 days under high NaCl conditions but not following prestimulation under low NaCl conditions (**Figure 5A**). We then cocultured T cells and MART-1 peptide-pulsed melanoma cells in the concomitant presence of high extracellular NaCl concentrations and observed a strong increase in melanoma cell lysis under high-NaCl conditions (**Figure 5B**). Given that we could rule out a direct effect of NaCl on melanoma cell growth or lysis **(Figure S6C, D)**, we attributed this finding to the effect of NaCl on antigen-specific CD8^+^ T cells. This ability of NaCl to promote CD8^+^ T-cell cytotoxicity was also corroborated with MART-1 specific T-cell clones, which we selected from the natural repertoire of healthy human HLA-A2-seropositive donors by tetramer staining **(Figure S6E, F)**.

**Figure 5.**
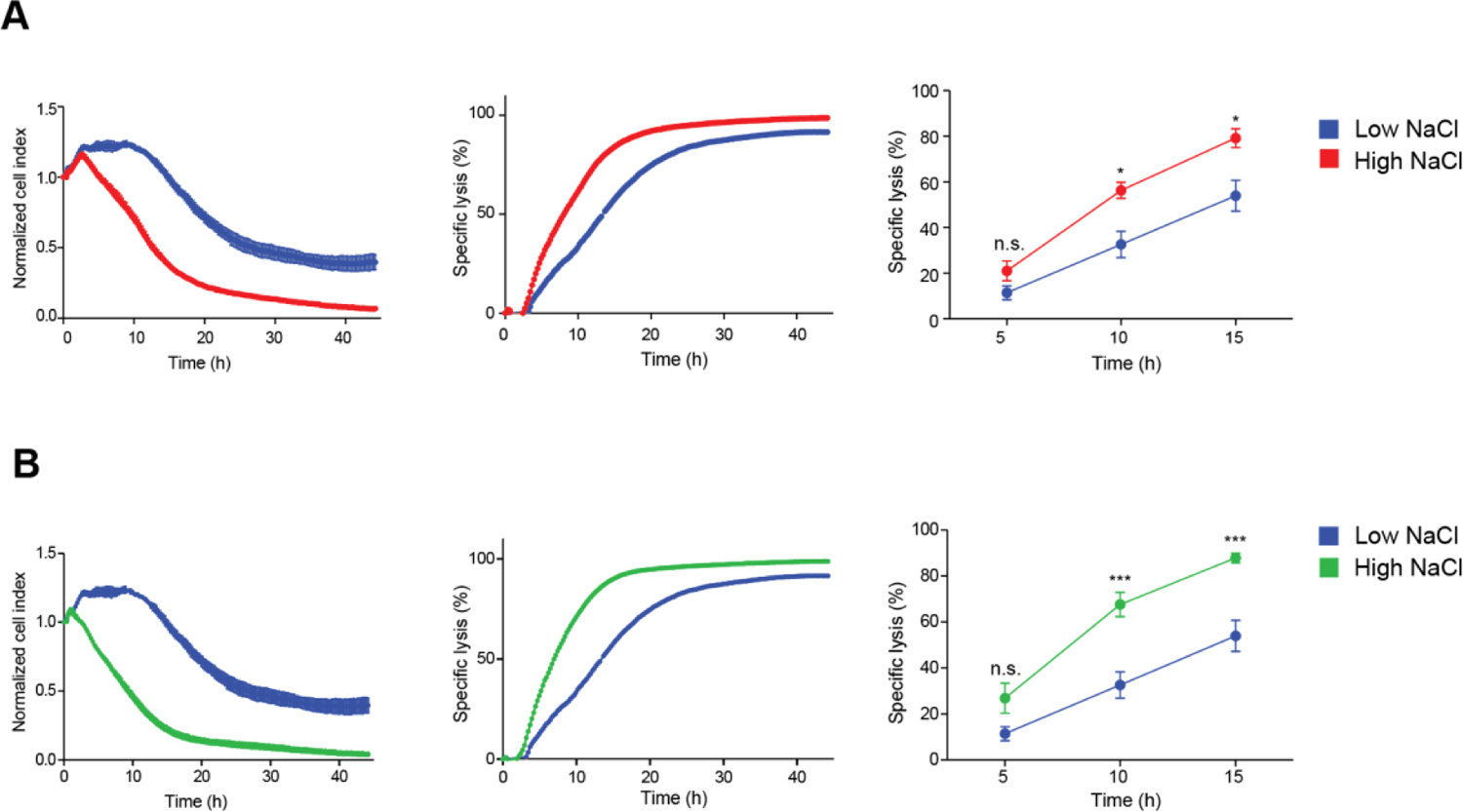
High NaCl conditions license human CD8^+^ T cells for superior killing of tumor cells. **(A, B)** Real time killing assay with nucleofected MART-1-specific T cells and A375 melanoma cell target cells at a 1:1 ratio under high and low NaCl conditions using the xCELLigence technology. n=3, mean ± S.E.M. Left, shown is the normalized cell index, middle, the specific lysis, right, the cumulative quantification of 3 T-cell donors. n=3, mean ± S.E.M. One-way ANOVA. **(A)** Prestimulation of MART-1-specific T cells under high and low NaCl conditions before coculture with MART-1 peptide pulsed target cells under low NaCl conditions. **(B)** Prestimulation as in (A) and coculture under high NaCl conditions.

Taken together, these findings demonstrate that the tumor antigen-specific cytotoxicity of human CD8^+^ T cells is enhanced in high NaCl, but not low NaCl, microenvironments. This enhanced killing ability could be stably maintained in CD8^+^ T cells prestimulated with high NaCl even after withdrawal from a high NaCl microenvironment.

### NaCl reduces tumor growth in a mouse model of pancreatic carcinoma

Given the enhanced cytotoxicity that was exerted by NaCl on T cells *in vitro,* we asked whether NaCl could be exploited for the enhancement of CD8^+^ T cell cytotoxicity during adoptive T cell therapies. We therefore investigated the impact of NaCl on CD8^+^ T-cell antitumor functions *in vivo.* To this end, we chose a well-established pancreatic cancer mouse model and used the pancreatic cancer cell line Panc02, which expresses the model antigen ovalbumin (Panc^Ova^), as described previously (Lutz et al., 2023). Naïve CD8^+^ T cells were obtained from OT-I mice, in which CD8^+^ T cells are specific for OVA_257-264_. These cells were stimulated *in vitro* with CD3 and CD28 mAbs under high (+30 mM) or low NaCl conditions for 3 days, washed and then adoptively transferred into mice with established Panc^OVA^ tumors. Seven days previously, the mice were injected subcutaneously (*s.c*) with 1 × 10^6^ Panc^OVA^ cells into the left flank (**Figure 6A**). Compared with untreated mice, mice treated with CD8^+^ T cells stimulated with low NaCl concentrations rejected tumors, as evidenced by reduced tumor volume (**Figure 6B**). Notably, tumor rejection was significantly increased following transfer of CD8^+^ T cells from high NaCl conditions (**Figure 6B**). NaCl did not affect CD8^+^ T-cell proliferation or viability *in vitro* (**Figure S7A, B**). Similar to the results observed in human CD8^+^ T cells, murine CD8^+^ T cells released significantly more granzyme B into the supernatant upon differentiation under high than under low NaCl conditions (**Figure 6C**), indicating that the increased tumor rejection was supported by enhanced release of this cytotoxic mediator. On day 3 of differentiation, murine CD8^+^ T cells also demonstrated enhanced intracellular granzyme B levels under high NaCl conditions as compared to low NaCl conditions, but there was no difference in the levels of other CD8^+^ T-cell effector cytokines, such as IFN-γ or TNF-α (**Figure 6D, Figure S7C, D**). Together, these data demonstrate that NaCl equips CD8^+^ T cells with improved antitumor functions *in vivo*.

**Figure 6.**
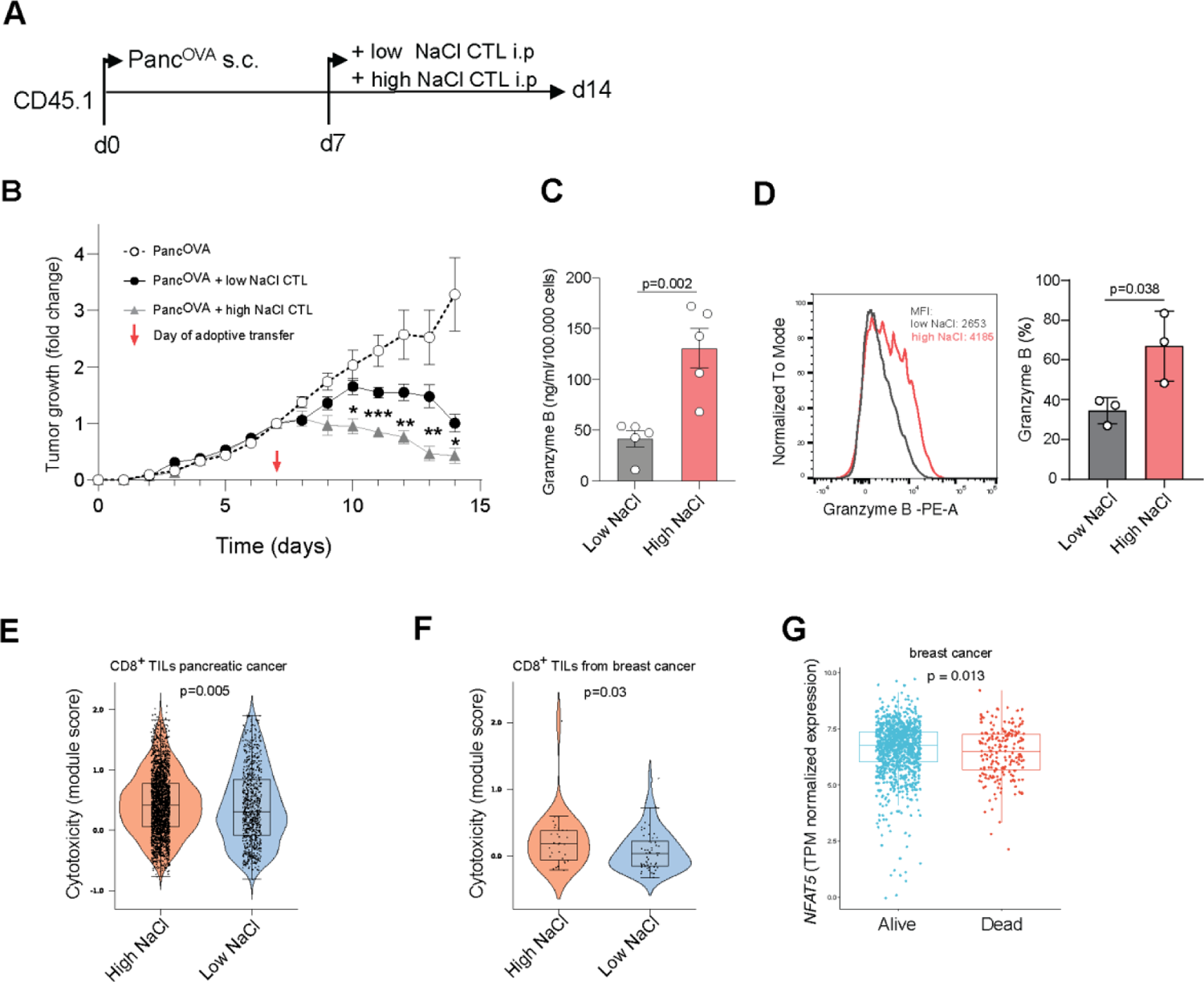
High NaCl conditions license adoptively transferred CD8^+^ T cells for enhanced tumor killing in a pancreatic cancer mouse model *in vivo*. (A) Experimental design. Panc^OVA^ tumor cells (1 × 10^6^) were subcutaneously injected into CD45.1 mice. CD8^+^ T cells from OT-1 mice were isolated from the spleen and lymph nodes and stimulated for 3 days under high (+30 mM) or low NaCl conditions before adoptive transfer of 0.5 × 10^6^ CD8^+^ T cells into tumor-bearing mice on day 7 after tumor cell injection. (B) The tumor growth curves of subcutaneous tumors are shown (mean ± S.E.M., n=6-7). Tumor growth was normalized to the tumor size on the day of CD8^+^ T-cell injection. Two-way ANOVA with Tukey’s HSD multiple comparison test. (C) ELISA with cell culture supernatants from CD8^+^ T cells stimulated for 3 days with CD3 and CD28 mAbs under high and low NaCl conditions. (D) Intracellular staining and flow cytometry on day 3 after stimulation with CD3 and CD28 mAbs for 3 days and PMA/ionomycin restimulation in the presence of brefeldin A. Paired t test (n=3). Left, representative experiment; right, cumulative quantification. Paired t test. All data points represent biological replicates. (E) scRNA-seq data of intratumoral cells from 51 patients with pancreatic cancer were obtained from GSE 155698, GSE111672, GSE154778, GSM4293555, PRJCA001063 (Chijimatsu et al., 2022) (integration of all cells: 10.5281/zenodo.6024273) and then filtered for CD8^+^ T cells as in Fig. 1E by marker gene expression. CD8^+^ T cells were categorized into cells with a high versus low NaCl signature based on the NaCl signature obtained from the single-cell transcriptomic comparison of CD8^+^ memory T cells treated with high vs. low NaCl concentrations as in Fig. 2C with cutoff defined as module score ≥ 0 and < 0 for high vs. low NaCl signature, respectively. Shown is the expression of cytotoxic gene expression defined as a module score obtained from published reports (*CTSW, GNLY, GZMA, GZMB, GZMB, GZMH, IFNG, KLRB1, KLRD1, KLRK1, NKG7, PRF1*) (Guo et al., 2018; Tirosh et al., 2016). Wilcoxon rank sum test was used to compute significance between the two subsets. The significance has been validated with genes from GO:0001916 (“positive regulation of T cell mediated cytotoxicity”) showing a significance with p=0.01. (F) scRNA-seq of intratumoral immune cells from patients with breast cancer (n=3) as in Fig. 1E filtered for CD8^+^ T cells by marker gene expression. The module score for the transcriptomic NaCl signature (obtained as described in Fig.1D) was used to distinguish CD8^+^ T cells with a high versus low NaCl signature with cutoff defined as module score ≥ 0 and < 0 for high vs. low NaCl signature, respectively. Shown is expression of cytotoxic gene expression defined and as a module score obtained from published reports as in (E) (Guo et al., 2018; Tirosh et al., 2016). Wilcoxon rank sum test was used to compute significance between the two subsets. The significance has been validated with genes from GO:0001916 showing a significance with p=0.05. (G) TPM normalized *NFAT5* expression in TCGA breast cancer patients grouped by the indicated metadata on survival. Wilcoxon rank sum test.

Finally, we tested whether T cell cytotoxicity was enhanced in the human tumor microenvironment in response to NaCl sensing. To this end, we interrogated the expression of the transcriptomic NaCl signature, which we had generated earlier, in tumor-infiltrating CD8^+^ T cells from 56 pancreatic cancer patients (10.5281/zenodo.6024273) (Chijimatsu et al., 2022). The pancreatic tumors displayed different amounts of tumor infiltrating CD8^+^ T cells as expected given the heterogeneity of this disease (**Figure S8A, B)**. Five tumors were excluded because they did not contain CD8^+^ T cells. Differential expression levels of the transcriptomic NaCl signature among individual TILs and their respective correlation with cytotoxicity was assessed. Only a subgroup of TILs within the tumor microenvironment was responding to NaCl (2511 TILs vs. 776 TILs). TILs within the subgroup of cells displaying a NaCl signature demonstrated significantly increased expression of cytotoxic gene signatures (**Figure 6E**). This correlation of a cellular NaCl signature with cytotoxicity supported the anti-tumor effect of NaCl within the complex human microenvironment. It further highlighted the differential impact of NaCl among TILs in the tumor microenvironment. To further validate these findings, we also revisited the tumor-infiltrating CD8^+^ T cells in the three breast cancer patients, for which we had earlier detected higher intratumoral than peritumoral NaCl concentrations (Azizi et al., 2018). Again, we found heterogeneity within the TILs from breast cancers with respect to NaCl sensing (34,5% TILs with positive NaCl signature) and a significant association of T cell cytotoxicity with expression of the transcriptomic NaCl signature (**Figure 6F**). We then also revisited the 1092 breast cancer patients from the TCGA source and categorized them into two groups based on the provided metadata, specifically into patients surviving and patients dying from cancer. Interestingly, we found that higher gene expression levels of *NFAT5*, which we and others have previously shown to be a NaCl-sensitive transcription factor in a great variety of cells (Matthias et al., 2019a; Miyakawa et al., 1999), was expressed at higher levels in the lesional tissue of breast cancer patients surviving breast cancer than in the lesional tissue of those patients dying from breast cancer (**Figure 6G**). Cumulatively, these results highlight the beneficial effect of NaCl within the tumor microenvironment for anti-tumor T cell immunity *in vivo* and for harnessing cytotoxicity in adoptive T cell therapies.

## Discussion

We have discovered that high NaCl concentrations in the extracellular microenvironment have a significant impact on cellular metabolism and enhance the effector functions of human CD8^+^ T cells. This leads to improved antitumor cytotoxicity *in vitro* and *in vivo*.

Potassium was previously proposed to regulate T-cell associated antitumor immunity (Eil et al., 2016). Potassium promotes both T-cell paralysis (Eil et al., 2016) and stemness with enhanced downstream T-cell effector functions (Vodnala et al., 2019). We found that sodium also acts as an ionic T-cell checkpoint and compartmentalizes in tissues in association with malignancy. We have discovered sodium to be a functionally active determinant of the tumor microenvironment with profound effects on the overall transcriptome of the tumor tissue and on CD8^+^ TILs, in particular. Sodium imposed a transcriptomic signature on human CD8^+^ memory T cells, which significantly differed from the signature of CD8^+^ T cells from low NaCl conditions. Remarkably, almost all tumor entities from the Cancer Genome Atlas demonstrated evidence of a transcriptomic sodium imprint as compared to healthy control tissues, pointing to sodium as a relevant, previously overlooked, constituent of the tumor microenvironment.

Despite overall higher intratumoral NaCl concentrations, the spatial distribution of NaCl within the tumor microenvironment, remains to be discerned. We suspect a differential NaCl distribution pattern within the tumor, which would be in line with our finding that T cells within the breast tumor microenvironment display heterogeneous expression of the transcriptomic NaCl signature, which corresponds to differences in their cytotoxicity levels. T cells in proximity to higher NaCl concentrations could therefore have an advantage for tumor control in an overall immunosuppressive cancer microenvironment. This raises the question as to the driving factors and mechanism of sodium deposition in tumors. Potassium has previously been shown to accumulate in tumor tissue as a result of cell necrosis and thus liberation from its intracellular compartment, where it promotes immune paralysis but also T cell stemness (Eil et al., 2016; Vodnala et al., 2019). However, intracellular sodium concentrations are low, suggesting that a different mechanism must underlie sodium enrichment in tumors. We have previously shown in different healthy human tissues, such as skin and muscle, that tissue sodium concentrations in humans are correlated with the level of glycosaminoglycan (GAG) deposition (Fischereder et al., 2017). These negatively charged macromolecules can locally trap cations such as Na^+^. Interestingly, GAGs have been reported to have increased abundance in tumor tissues (Theocharis and Karamanos, 2019; Wei et al., 2020). Additionally, the osmosensitive transcription factor NFAT5, which acts upstream of SGK-1, has previously been described to regulate GlcAT-1, a key enzyme for GAG biosynthesis (Hiyama et al., 2009). Sodium accumulation in the tumor microenvironment could therefore reflect a bystander effect from GAG deposition. Its molecular regulation could offer potential therapeutic targets for tumor immunity, given the profound effects of sodium on cytotoxic CD8^+^ T-cell effector functions and possibly cancer survival. Moreover, this intratumoral sodium deposition could also serve diagnostic purposes in cancer as already suggested by ^23^Na^+^ MRI studies (Zaric et al., 2016). Other mechanisms of sodium accumulation in human tumors, including dietary influences, as previously suggested in grafted tumor mouse models, could also contribute to local sodium accumulation and remain to be investigated in the future (He et al., 2020).

An intriguing observation of this study was the metabolic burst in human CD8^+^ T cells exposed to high NaCl concentrations, which was associated with potentiated effector functions and, ultimately, tumor cell killing. This finding was unexpected, considering that T cells in tumor microenvironments are often found to be paralyzed due to chronic stimulation and exposure to immunosuppressive signals, such as potassium or TGF-β, from the microenvironment (Batlle and Massague, 2019; Eil et al., 2016). However, our specific analysis of activation parameters, effector functions, cellular bioenergetics and single-cell transcriptomic trajectories confirmed the enhanced effector identity of CD8^+^ T cells in response to high NaCl concentrations at the transcriptomic, proteomic, metabolomic and functional levels. NaCl and its previously reported downstream signaling targets, such as NFAT5 and SGK-1, therefore represent attractive antagonistic players to be harnessed in an overall immunosuppressive tumor microenvironment for acute bursts of cytotoxicity. Stemness, a relevant cellular parameter for the maintenance of antigen-specific tumor immunity, which is known to be inversely correlated with effector differentiation (Vodnala et al., 2019), was, in accordance with the enhanced effector functionalities, reduced under high NaCl conditions in CD8^+^ T cells. This stresses the fact that competitive signals from a complex microenvironment act on tumor-infiltrating lymphocytes. The differential integration of the various tissue signals, which are overall considered to be immunosuppressive in cancer microenvironments, will ultimately determine the overall outcome in terms of cancer progression. Additionally, the effects of NaCl on other cellular players, such as myeloid-derived suppressor cells, whose expansion has recently been found to be suppressed by NaCl, or on Th17 cells, need to be considered in the net arithmetic evaluation of antitumor immunity (He et al., 2020; Kleinewietfeld et al., 2013; Wu et al., 2013). Our finding that surviving patients in a large cohort of breast cancer patients (>1000, TCGA) display increased intratumoral expression of the sodium sensor *NFAT5* compared to patients who died from breast cancer, already points to an advantageous clinical outcome in association with increased cellular salt sensing.

Our results showing the enhanced cellular metabolism of NaCl-conditioned CD8^+^ T cells are in stark contrast to the recently reported perturbation of mitochondrial respiration in Treg cells upon exposure to high NaCl concentrations (Corte-Real et al., 2023). While CD8^+^ T cells generated increased ATP via engagement of multiple layers of cellular metabolism, high NaCl was reported to decrease the metabolic fitness (Macintyre et al., 2014) of Treg cells through inhibitory effects on mitochondrial respiration (Corte-Real et al., 2023). As a consequence, Treg cells lose their suppressive functions and acquire the ability to produce proinflammatory cytokines such as IFN-γ. These opposing effects of NaCl on the cellular metabolism of both T cell lineages could be due to previously well characterized intrinsic differences in their baseline metabolic identity, with Treg cells depending on lipid oxidation and conventional T cells depending on glycolytic activity to exert their assigned functions (Buck et al., 2015). This distinct metabolic wiring of CD8^+^ T cells in response to NaCl also translates into a differential dependence on Glut-1, the glucose importer, which we found to be strongly upregulated by NaCl in CD8^+^ memory T cells (Macintyre et al., 2014). Notably, Glut-1 contains a consensus site ((95)Ser) for phosphorylation by SGK1 (Palmada et al., 2006), the intracellular sensor for NaCl, which we also found to be upregulated in a transcriptomic comparison of CD8^+^ T cells in high NaCl conditions. This finding argues for a direct mechanistic link between NaCl-sensing and glucose metabolism. Additionally, experimental differences, such as short versus sustained NaCl exposure in Treg and CD8^+^ T cells, respectively, could account for the differential impact of NaCl on these different cells (Corte-Real et al., 2023; Pearce et al., 2013). Notably, both the loss of Treg suppressive functions and the potentiation of CD8^+^ T cell cytotoxicity, are known to jointly contribute to improved tumor elimination (Shan et al., 2022). Although the impact of NaCl for tumor control by Treg cells has not been formally investigated thus far, we speculate that NaCl enrichment in the tumor microenvironment could therefore serve to pull together different cellular functions to support an improved overall antitumor response despite alternative cellular engagement of metabolic pathways in distinct immune cell types.

The significant reduction in tumor size upon adoptive transfer of CTLs stimulated under high NaCl conditions in the tumor mouse model of pancreatic cancer was compelling, given the overall poor outcome of solid cancers and of pancreatic carcinoma, in particular (Hidalgo, 2010). We chose an adoptive T-cell transfer model to pinpoint NaCl-induced immunomodulation of T cells and to exclude potential systemic effects of NaCl on the microbiota, on the malignant cells themselves or on the immune surveillance of other cell types. Our *in vivo* data suggest an easy and safe actionable strategy to dramatically increase T-cell cytotoxicity via metabolic regulation, with beneficial effects for cancer and possibly chronic infections. It is now tempting to speculate that preconditioning CAR-T cells with elevated NaCl concentrations would result in improved antitumor responses in patients. Increased T cell effector functions, however, come at the expense of stemness and thus long-term maintenance of tumor-antigen or pathogen-specific memory T cells (Vodnala et al., 2019). NaCl therefore assumes the role of a metabolic switch factor at the interface of T cell effectorness and stemness. It should also be noted that the continued NaCl deposition in tumors could favor T cell exhaustion in the long run, especially in slowly growing “old” solid tumors.

Our finding that TILs with a higher NaCl signature display higher cytotoxicity and the finding that tumors with higher expression of the NaCl-sensing transcription factor NFAT5 correlate with survival in breast cancer patients support the beneficial effect of NaCl on a subset of TILs in an otherwise complex and overall immunosuppressive microenvironment. Clinical studies will have to reveal in the future whether ionic conditioning of T cell products with NaCl will lead to beneficial outcomes for cancer patients. In summary, these findings advance our understanding of CD8^+^ T cell regulation and may lead to new therapeutic strategies for the treatment of cancer and other pathologies that may benefit from augmented T-cell cytotoxicity.

## Methods

### Cell purification and sorting

Peripheral blood mononuclear cells (PBMCs) from fresh peripheral blood of healthy donors were isolated by density gradient centrifugation using Ficoll-Paque Plus (GE Healthcare). Cytotoxic T cells and T helper cells were isolated from the PBMCs by positive selection with CD8- and CD4-specific MicroBeads (Miltenyi Biotec), respectively, using an autoMACS Pro Separator (Miltenyi Biotec). Memory T cells were isolated as CD45RA^-^ lymphocytes, and naïve T cells were isolated as CD45RA^+^CD45RO^-^ CCR7^+^ lymphocytes with a purity of over 98%. The antibodies used for flow cytometry cell sorting were described previously (Noster et al., 2016; Zielinski et al., 2012). The cells were sorted with a BD FACSAria III (BD Biosciences), BD FACSAria Fusion (BD Biosciences), or a Cytek Aurora CS (Cytek).

### Cell culture

Human T cells were cultured in RPMI-1640 medium supplemented with 2 mM glutamine, 1% (vol/vol) nonessential amino acids, 1% (vol/vol) sodium pyruvate, penicillin (50 U/ml), streptomycin (50 μg/ml; all from Invitrogen) and 10% (vol/vol) fetal calf serum (FCS; Biochrom). For antigen-specific assays, FCS was replaced with 5% human serum (Sigma-Aldrich, H6914). Hypersalinity (high NaCl) was induced by increasing the NaCl concentration to 185 mM as described previously unless indicated otherwise (Matthias et al., 2020; Matthias et al., 2019a). The final sodium concentrations in the cell culture media were confirmed by potentiometry with a Cobas 8000 analyzer (Roche). T cells were stimulated with plate-bound CD3 (1 μg/ml, clone TR66) and CD28 mAbs (1 μg/ml CD28.2; BD Biosciences). T-cell clones were generated under nonpolarizing conditions, as described previously, following single-cell deposition via flow cytometry-assisted cell sorting or by limiting dilution plating (Noster et al., 2014).

### Generation and isolation of antigen-specific T cells

To obtain tumor antigen-specific T cells (MART-1 specificity), 1 x 10^7^ freshly isolated PBMCs were washed twice in PBS and resuspended in preequilibrated nucleofection buffer 1SM (Chicaybam et al., 2013). Twenty micrograms of transposon coding MART-1-specific TCR/0.5 µg of transposase SB100X solution was added to the cell suspension, which was subsequently transferred into the electrophoresis chamber. Nucleofection was performed on a Lonza Nucleofector IIb. After nucleofection, the cell suspension was transferred into a 24-well microplate containing complete culture medium with human serum and 50 U/ml IL-2. After 24 hours, MART-1-specific T cells were isolated from nucleofected PBMCs with a BD FACSAria III cell sorter (BD Biosciences) by selecting CD3-FITC and anti-mouse-TCR β-chain-APC positive cells after exclusion of dead cells with propidium iodide.

MART-1-specific CD8^+^ T cells from the natural T-cell repertoire were isolated from PBMCs from an HLA-A2-seropositive healthy donor after tetramer staining in MACS buffer (PBS supplemented with 1 % (vol/vol) fetal calf serum, 2mM EDTA) at 4 °C for 30 minutes. Tetramers were assembled the day before by incubating the pMHC complex with streptavidin-APC or with streptavidin-BV421 overnight at 4 °C. The tetramers used for staining were pMHC-Streptavidin-BV421 and pMHC-streptavidin-APC. The antibodies used for staining were CD3-FITC, CD8-PE and CD19-ECD. To exclude dead cells, propidium iodide was used. Single-cell sorting was carried out at a MoFlo Astrios Cell sorter (Becton Dickinson) after gating on single CD3^+^CD8^+^CD19^-^PI^-^ pMHC-Streptavidin-APC^+^/pMHC-Streptavidin-BV421^+^ lymphocytes. pMHC complexes loaded with MART-1 peptide were refolded as described previously (Altman et al., 1996) (Busch et al., 1998) (Garboczi et al., 1996a) (Garboczi et al., 1996b). Antigen-specific T-cell clones were generated as described previously (Braun and Zielinski, 2014; Zielinski et al., 2012).

### Cytotoxicity assay

The viability of A375 melanoma cells (target cells) was determined in real time using an xCELLigence SP Real-Time Cell Analyzer (ACEA Biosciences), which allowed the quantitative and continuous monitoring of adherent A375 melanoma cells (target cells) through the measurement of electrical impedance every 15-30 minutes. For baseline measurements, 100 µl of A375 growth medium was added to the 96-well E-plate. A375 melanoma cells were seeded onto the E-plate at a density of 5 x 10^3^ cells/100 µl of growth medium. After 24 hours, 100 µl of culture medium was replaced by 10^-7^ M MART-1 peptide in 100 µl of fresh medium. After 1 h of incubation with the peptide, MART-1 specific CD8^+^ T cells, which had been preactivated in high or low NaCl conditions for 5 days (or no T cells as a control), were added at a 1:1 ratio in 100 µl after removal of an equal amount of the growth medium. All conditions were set up in technical duplicates. Cell indices were monitored every 15-30 minutes for another 24-48 hours on the xCELLigence System at 37 °C with 5 % CO_2_. Values are represented as cell indices (CI), cell indices normalized to the start of the co-culture (nCI) or as cell lysis calculated according to the following formula: 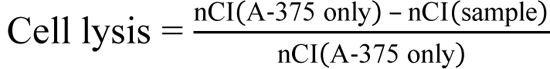.

### Flow cytometric analysis of cytokines, transcription factors and surface markers

For intracellular cytokine staining, human cells were restimulated for 5 h with PMA and ionomycin with brefeldin A being added for the final 2.5 h of culture (all Sigma–Aldrich). The cells were fixed and permeabilized with Cytofix/Cytoperm (BD Biosciences for cytokines, eBioscience for transcription factors) according to the manufacturer’s instructions. The cells were stained with anti-cytokine antibodies and were analyzed with a BD LSRFortessa (BD Biosciences), a CytoFLEX (Beckman Coulter), MACSQuant Analyzer (Miltenyi Biotec) or a Cytek Aurora Analyzer (Cytek). Flow cytometry data were analyzed with FlowJo software (Tree Star) or with Cytobank (Cytobank Inc.). Cytokines in culture supernatants were quantified by ELISA (R&D Systems) or by Luminex (Thermo Fisher Scientific) according to standard protocols. High-dimensional cell surface phenotyping of sorted human memory CD8^+^ T cells from 4 individual healthy blood donors was performed with a newly established 33-colour panel for surface markers on a Cytek Aurora Analyzer. The cells were analyzed on day 5 after stimulation with CD3 and CD28 mAbs under high and low NaCl conditions. Inverse hyperbolic transformation was performed on the indicated fluorescence markers using the “arcsinhTransform” and “transform” functions of the flowCore (v.2.6.0) R package (R v.4.1.3) (Hahne et al., 2009). All transformed samples were subjected to quality control and filtered using “PeacoQC” (v.1.8.0) based on both the Median Absolute Deviation and the IsolationTree (Emmaneel et al., 2022). Each sample was then randomly downsampled to 1420 cells using the base R package to match the lowest cell population number after preprocessing. The resulting expression matrices were concatenated and analyzed using the “DetermineNumberOfClusters” function of FlowSOM (v.2.2.0) using the “metaClustering_consensus” method (Van Gassen et al., 2015) to determine the number of metaclusters per blood sample (median of 22 metaclusters). Using the default 10 × 10 grid, 100 phenotypic clusters were generated and were grouped into the 22 metaclusters using FlowSOM.

The comparison of clusters between high and low salt conditions was performed using the “GroupStats” function of FlowSOM. Significant differences within clusters were defined according to a fold change computed using the Wilcoxon test of over 2.5 and under −2.5. The minimal distance trees were plotted using the “PlotStars” function of FlowSOM for both conditions separately sizing the nodes according to the median fluorescence intensity (MFI) per group.

### Ca^2+^ flux measurement

T cells were collected at day 5 after prestimulation with CD3 and CD28 mAbs under low and high NaCl conditions. A total of 5 × 10^6^ cells were resuspended in 500 μl of cell culture medium with 5 μg/ml of Indo-1 AM and 0.5 μg/ ml of pluronic F-127 (both from Life Technologies) and incubated in the dark for 30 min at 37 °C. The cells were washed, then resuspended in 500 μl cell culture medium and kept on ice until measurement. For each Ca^2+^ flux measurement, cells were diluted with 37°C pre-warmed cell culture medium and analyzed with a MaxQuantX Flow Cytometer (Miltenyi Biotech) or BD FACSAria Fusion Flow cytometer (BD Biosciences). After acquisition of the baseline fluorescence level, the TCR was stimulated by addition of 15 μg/ml of the CD3 antibody clone UCHT1 or alternatively by restimulation with CD3 mAbs (TR66 clone) and crosslinking by 1.3 μg/mL anti-mouse IgGF(ab)’_2_ fragments (Jackson ImmunoResearch). The ratio of Ca^2+^-bound Indo-1 versus Ca^2+^-free Indo-1 was analyzed using FlowJo 9 software (Beckman Coulter).

### Metabolic assays

The mitochondrial function (oxidative phosphorylation) and glycolytic rate of CD8^+^ memory T cells, which were prestimulated for 5 days with CD3 and CD28 mAbs under high versus low NaCl conditions, were assessed after washing using a Seahorse XFp Analyzer (Agilent Technologies, Waldbronn, Germany). T cells were resuspended in Seahorse assay medium (pH adjusted to 7.4 – 7.45) on XF96 cell culture microplates at a density of 2.5 x 10^5^ cells per well. Cells were centrifuged for 5 min at 400x g to adhere and form a monolayer at the bottom of the plate. All experiments were performed with 4 technical replicates. The XF^e^96 extracellular flux assay kit (Agilent Technologies, Waldbronn, Germany) was used according to the manufacturer’s protocol with addition of oligomycin, CCCP, antimycin A and rotenone at concentrations of 2 x 10^-6^ M, 1,5 x 10^-6^ M, 2 x 10^-6^ M and 2 x 10^-6^ M, respectively. Evaluation and calculation of mitochondrial and glycolytic indices was done with the Wave software 2.2.0 (Agilent Technologies, Waldbronn, Germany). Glut-1 expression by T cells was analyzed by flow cytometry using anti-hGlut1-PE (R&D, FAB1418P) or anti-hGlut-1-FITC (R&D, FAB1418F). Fatty acid uptake was analyzed by flow cytometry using Bodipy (Invitrogen, Bodipy FL C16, 1 μM, Excitation/Emission 505/512 nm on FITC channel).

### Gene expression analysis

For mRNA-seq analyses (bulk transcriptome), CD8^+^ memory T cells were isolated *ex vivo* as described above and stimulated for 48 h with CD3 and CD28 mAbs for a total culture time of 5 days in the presence of low and high NaCl concentrations (GSE232365, password encoded until day of publication). Total RNA was extracted from cells lysed in TRI reagent (Sigma-Aldrich) according to the manufacturer’s protocol. RNA was quantified using a NanoDrop 2000 spectrophotometer (Thermo Fisher Scientific) and its quality was verified by an Agilent 2100 Bioanalyzer (Agilent) according to the manufacturer’s guidelines.

Library preparation for RNA-seq was performed using the TruSeq® Stranded Total RNA Sample Preparation Guide (Illumina). The barcoded libraries were sequenced on a NovaSeq 6000 platform (Illumina) by Novogene (Cambridge, UK) with paired-end, 150-bp reads (PE150). Approximately 2 Gb of sequencing reads were produced on average per sample. The reads were mapped to the reference transcriptome built from the human genome assembly hg38 (GRCh38) using STAR v2.6.1a (Dobin et al., 2013). Transcripts were quantified with salmon v0.11.3 in the alignment-based mode (Patro et al., 2017).

Downstream analyses were performed with the statistical framework R (Team, 2022). Significantly differentially expressed genes were identified using the R package DESeq2 (Love et al., 2014), using a false discovery rate (FDR) corrected significance threshold 0f 0.05. Log fold change cutoff was set at 0 for determining differentially expressed genes using the Wald Test and the significance threshold was based on the false discovery rate (FDR) < 0.05. Of 60623 tested genes we identified 26837 with a non-zero mean expression. Of these, 1956 were significantly up- and 1926 down-regulated. Plots were produced with the R package ggplot2 (Wickham, 2016). The R package clusterprofiler v4.6.0 ^23^ was employed for overrepresentation analysis as well as gene set enrichment analysis and visualization of GO terms within the ontology “Biological Process” and KEGG pathways. The enrichment plots show the top 20 entries for the significantly up-regulated, down-regulated and dysregulated genes respectively, which were ranked according to the shrunken fold change values calculated by DESeq2, as previously described (Love et al., 2014). Barcode plots of enriched gene signatures were generated with clusterProfiler. Heatmaps were generated using the R package ComplexHeatmap v2.14.0 (Gu et al., 2016). Overlay of expression data over KEGG pathways was done with the R package pathview v1.38.0 (Luo and Brouwer, 2013).

### Transcriptomic profiling of human cancers from the Cancer Genome Atlas (TCGA)

Gene expression data of patients with cancer were obtained from The Cancer Genome Atlas (TCGA). Non-diseased tissue-specific expression data were collected from the Genotype-Tissue Expression (GTEx) project. Gene expression data from both projects were re-preprocessed by the same RNA-seq pipeline from the UCSC RNA-seq Compendium (Goldman et al., 2020).. The resulting gene expression data was made publicly available as TCGA-TARGET-GTEx cohort (https://xenabrowser.net/datapages/?cohort=TCGA%20TARGET%20GTEx), which was used for subsequent transcriptomic profiling of human cancers. Data from the TARGET cohort (pediatric data, https://www.cancer.gov/ccg/research/genome-sequencing/target) was discarded. We downloaded “Transcripts Per Kilobase Million” (TPM) normalized RNA sequencing based expression data for 60498 genes from the Xena platform as well as associated metadata (https://toil.xenahubs.net/download/TcgaTargetGtex_rsem_gene_tpm.gz, https://toil.xenahubs.net/download/TcgaTargetGTEX_phenotype.txt.gz, downloaded March 6, 2023)^1^. By using data from the toil-hub, it was ensured that data was recomputed with one pipeline, thus circumventing batch effects introduced by different pipelines. After filtering for tumor entities that included healthy control tissues, 8972 “Primary Tumor” samples and 727 “Solid Tissue Normal” samples of 9019 patients (TCGA dataset) and 4472 “Normal Tissue” samples of 535 donors (GTEx dataset) across 25 tumor types were available. Expression data for healthy samples (“Solid Tissue Normal”, “Normal Tissue”) were pooled per tumor site. To enable gene set enrichment analysis (GSEA), log_2_(fold-changes) of gene expression, data were calculated per gene and tumor site between mean TPM-expression data of tumor and healthy samples. Subsequent GSEA using all 1956 up-regulated genes of the salt signature and the log_2_(fold-changes) ranked gene list per tumor site as input was performed using the fgsea package (v1.2.4.0) within the R statistical programming environment (v4.2.3) to compute NES and one-tailed test based p-values for positive enrichment of the gene signature. Default parameters were used for calculating GSEA with the function fgsea::fgsea except for minSize (=10) and maxSize (=4000) and scoreType (=”pos”).

TPM normalized expression levels of *NFAT5* expression per patient from the breast cancer patient cohort of the TCGA dataset were grouped according to survival status (dead or alive) that was provided by the TCGA metadata information. A Wilcoxon rank sum test was used for comparison of the dead versus alive patient groups.

### Single-cell mRNA sequencing analysis (scRNA-seq)

For scRNA-seq analysis of matched tumoral and peritumoral CD8^+^ T cells from patients with breast cancer, tissue samples from three patients with breast cancer (BC01, BC02, BC03) were reanalyzed (GSE114727) (Azizi et al., 2018). Quality control with removal of cells expressing fewer than 200 genes and of genes present in less than 3 cells was performed using the python package Scanpy v.1.9.1. Doublets were removed with Scrublet v.0.2.3. Data were normalized to 10,000 reads per cell using normalize_per_cell() and log-transformed using log1p() functions. Sample integration and batch correction were performed using the combat() function provided by Scanpy. CD8^+^ T cells were annotated with CellTypist v.1.3.0 using the pretrained model Immune_All_High.pkl. Cells assigned to a probability of being a T-cell < 0.6 were filtered out. Then, T cells with expression values of CD8A > 2.5 and CD8B > 2 were identified as CD8^+^ T cells. Transcriptomic NaCl signatures were derived from either a bulk transcriptomic comparison of FACS-sorted human CD8^+^ CD45RA^-^ T cells stimulated with CD3 and CD28 mAbs for 5 days under high versus low NaCl conditions (FDR < 0.05) or scRNA-seq of human CD8^+^ CD45RA^-^ T cells stimulated with CD3 and CD28 mAbs for 3 days under high versus low NaCl conditions (adjusted p value < 0.05). Significantly up- and downregulated gene sets from the two transcriptomic analyses were tested separately on the tumoral and peritumoral CD8^+^ T cells from the scRNA-seq analysis of the three patients with breast cancer. For GSEA, genes were ranked according to their expression values using the rank_genes_groups() function provided by Scanpy and analyzed with gseapy v.1.0.2. Module scores were calculated with the Scanpy score_genes() function. For comparisons of tumoral versus peritumoral CD8^+^ T cells, statistical significance was determined with the Wilcoxon rank sum test with the alternative hypothesis = ‘greater’ for upregulated gene sets and ‘less’ for downregulated gene sets.

For scRNA-seq (GSE232149, currently password encoded until publication), a library of human CD8^+^CD45RA^-^CD45RO^+^ cells that were stimulated with CD3 and CD28 mAbs for 3 days under high and low NaCl conditions was constructed with Chromium Next GEM Single Cell 5′ Reagents v.2 (Dual Index) (10x Genomics, Inc.). The library was sequenced on an Illumina NovaSeq 6000 Sequencing System (Flow Cell Type S4) according to the manufacturer’s instructions, with 150-bp, paired-end, dual-indexing sequencing (sequencing depth: 20,000 read pairs per cell). Read alignment and gene counting of the single-cell datasets were performed with CellRanger v. 7.0.1 (10x Genomics, Inc.). For downstream analysis, the filtered barcode matrix (CellRanger multi pipeline) was processed with the R package Seurat v.4.0.4. For quality control, cells with unique feature counts > 9000 and a count value > 80000 were filtered out. The total counts were normalized to 10,000 reads per cell. Each gene was centered and scaled to unit variance. Doublets were removed using the R package DoubletFinder v.2.0.3. UMAPs were generated using the RunUMAP() function and Leiden clustering was performed using the FindNeighbors() and FindClusters() functions from Seurat. The top 10 marker genes per Leiden cluster were determined with the wilcoxauc() function in the R package presto (v.1.0.0) and depicted in a heatmap using the R package ComplexHeatmap (v.2.12.1). Differential gene expression was evaluated using FindMarkers(), and genes were declared as significant based on an adjusted p value threshold of 0.05. Gene set enrichment analysis for Gene Ontology (GO) was conducted using clusterProfiler (v.4.4.4). Multiple testing correction was performed using the Benjamini-Hochberg method, and significantly enriched GO terms were visualized in a dotplot with clusterProfiler (v.4.4.4).

In analyses represented by violin plots, the python package Scanpy v.1.9.1 was used as an alternative to the R package Seurat v.4.0.4. Doublets were predicted and removed using Scrublet v.0.2.3 as before. Cells expressing fewer than 200 genes were filtered out. Cells were normalized to 10,000 reads per cell using normalize_per_cell() and log-transformed using log1p() functions. Gene expression values were scaled to a maximum value of 10. For UMAP, highly variable genes were computed using the function scanpy.pp.highly___variable___genes(). Principle Component Analysis (PCA) was performed using the function pca() n=30. Finally, the neighbors of each cell and the UMAP were computed using the functions scanpy.pp.neighbors() and scanpy.pp.umap(), respectively. Module scores were computed using the function score_genes() provided in the python package Scanpy v.1.9.1.

For trajectory analyses, diffusion pseudotime was used (Haghverdi et al., 2016). The cell with the highest expression of the stemness marker gene *TCF7* was chosen as the starting point for pseudotime inference. For validation, the analysis was repeated using the cell with the highest expression of the stemness gene set from a public data set (Wu et al., 2016). The function diffmap() and dpt() from Scanpy were used to compute the diffusion map representation and to assign pseudotime values to each cell in the dataset. For RNA velocity, the velocyto pipeline v.0.17.17 was used to obtain the pre-mature (unspliced) and mature (spliced) transcript information based on Cell Ranger output. The functions scv.pp.moments(), scv.tl.velocity() and scv.tl.velocity_graph() from scVelo v.0.2.5 were applied to recover RNA velocity and scv.pl.velocity_embedding_stream() was used for visualization. Quality control and preprocessing were performed using the Python package Scanpy v.1.9.1. Genes with a minimum count of 1 were retained and doublets were removed using Scrublet v.0.2.3. Total-count normalization was applied using normalize_total(). Data was log-transformed using log1p() and highly variable genes were identified using highly_variable_genes().

### Ion quantification by inductively coupled plasma optical emission spectroscopy (ICP-OES)

For sample preparation, frozen specimens were subject to a freeze dryer (Heraeus Christ, Osterode, Germany) and lyophilized at a – 35 °C condenser temperature until the weight became constant. Subsequently, the dried samples were transferred into closed quartz vessels and digested with HNO_3_ (suprapure, subboiling distilled; Merck, Darmstadt, Germany) in a Discover® SP-D 80 microwave digestion system (CEM corporation, Charlotte, North Carolina, USA). The resulting solution was brought to exactly to 10 mL with Milli-Q H_2_O and was then ready for element determination.

Element determination of sodium (Na) and potassium (K) was performed with inductively coupled plasma-optical emission spectrometry (ICP–OES) “ARCOS (Ametek-Spectro, Kleve, Germany). The measured spectral element lines in nm were K: 766.491 nm and Na: 589.592 nm. Sample introduction was carried out by a peristaltic pump, connected to a MicroMist nebulizer with a cyclone spray chamber. The RF power was set to 1400 W, the plasma gas was 15 L Ar /min, and the nebulizer gas was 0.6 L Ar /min. For quality control, three blank determinations and a control determination of a certified standard (CPI) for all mentioned elements were performed regularly after ten measurements. Analysis of the results was carried out on a computerized laboratory data management system, which related the sample measurements to calibration curves, blank determinations and control standards.

### Mouse experiments

Congenic C57BL/6 CD45.2^-^CD45.1^+^ mice were bred in-house (Animal Facility Philipps-University Marburg, BMFZ). OT-1 (B6.Cg-Tg(TcraTcrb)1100Mjb) mice were obtained from Jackson Laboratories. CD8^+^ T cells were obtained from the lymph nodes (LNs) and spleens of OT-1 mice using a negative selection kit (130-104-075, Miltenyi Biotec). For CTL differentiation, naïve CD8^+^ T cells were cultured in RPMI (10% FCS) and stimulated with plate-bound mCD3 mAbs (3 µg ml^-1^, clone 145-2C11, Biolegend), soluble mCD28 (0.5 µg ml^-1^, clone 37.51, Biolegend), rhIL-2 (50 Uml^-1^, Novartis) and anti-mIFN-γ (5 µg ml^-1^, clone XMG1.2, Biolegend) in the presence or absence of 30 mM NaCl. For intracellular cytokine staining, cells were restimulated after 72 h of culture with PMA (50 ng ml^-1^) and ionomycin (1 µg ml^-1^, both from Sigma-Aldrich) in the presence of brefeldin A (5 µg ml^-1^; Biolegend) for 4 h. Cells were fixed with 2 % formaldehyde for 20 min at room temperature. After fixation, intracellular staining for TNF-α-FITC (eBioscience), granzyme B-PE (eBioscience) and IFN-γ-APC (BioLegend) was performed in saponin buffer (0.1% saponin, 1% BSA in PBS) for 30 min at 4 °C. For proliferation, CD8^+^ T cells were labeled with the cell proliferation dye fluor670 (2 µM) for 10 min at 37 °C protected from light. The, cells were then washed, resuspended in culture medium and seeded into a 96-well-plate. Proliferation was measured by flow cytometry on day 3. Apoptosis was quantified using annexin V-propidium iodide (PI) staining. Cells were stained for 20 min in BSS with annexin V (APC #640920 Biolegend) at room temperature in the dark. PI (Invitrogen) was added immediately before flow cytometric analysis on an Attune NxT Cytometer (Thermo Fisher Scientific).

For *in vivo* animal experiments, 8–12-week-old CD45.1 mice were injected s.c. with 1 × 10^6^ Panc^OVA^ cells *s.c*. into the left flank. Seven days post-injection, tumor-bearing animals were treated with 0.5 × 10^6^ CTLs, which had been differentiated from naïve OT-1 CD8^+^ T cells for 3 days *in vitro* in the absence or presence of additional 30 mM NaCl. Tumor growth was measured, and tumor volume was calculated (V=length x width^2^ x 0.5; mm^3^).

Adherent Panc^OVA^ pancreatic adenocarcinoma cells (derived from Panc02) were grown in T75 flasks (Sarstedt) with DMEM (10% FCS), and split every 2-4 days upon reaching 70 % confluence, harvested by trypsinization (1x trypsin/EDTA, Sigma T-4174) and reseeded (5 x 10^5^ cells/10 ml medium in a T75 flask)). For selection, 500 mg/L G418 (G8168 Sigma–Aldrich) was added.

### Statistics

Statistical tests are indicated in the corresponding figure legends. Statistical tests for transcriptomic or metabolomic analyses are described in the corresponding sections. Error bars indicate the standard error of the mean (S.E.M.) unless otherwise stated; *p* values of 0.05 or less were considered significant. Analyses were performed using GraphPad Prism v.7-9.

### Study approval

Ethical approval was obtained from the Institutional Review Board of the Technical University of Munich (195/15s, 146/17s, 491/16s), the Charité-Universitätsmedizin Berlin (EA1/221/11) and the Friedrich Schiller University Jena (2020-1984_1). All work was carried out in accordance with the Declaration of Helsinki for experiments involving humans and with the Regierungspräsidium Giessen for studies involving mice.

## Supplementary Figures

**Figure S1.**
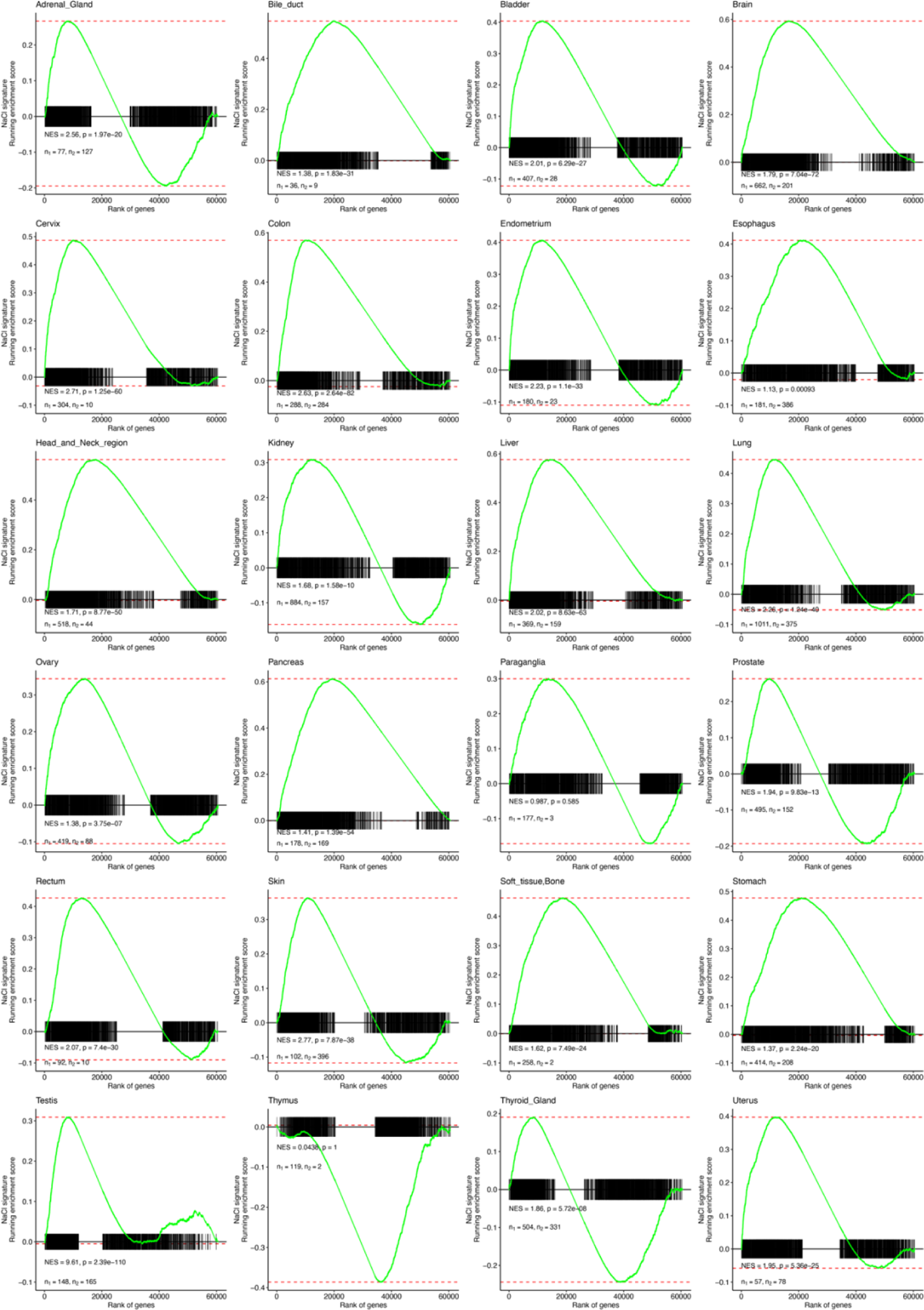
Intratumoral tissues from a wide range of solid tumors displays a transcriptomic enrichment of a NaCl signature. Barcode plots of GSEA results according to tumor tissue from The Cancer Genome Atlas (TCGA). Running enrichment score and sorted position of the 1956 significantly upregulated genes in the salt signature are shown. NES: normalized enrichment score, p: significance of the enrichment (one-tailed test for positive enrichment), n_1_: patients providing tumor samples, n_2_: patients providing healthy samples.

**Figure S2.**
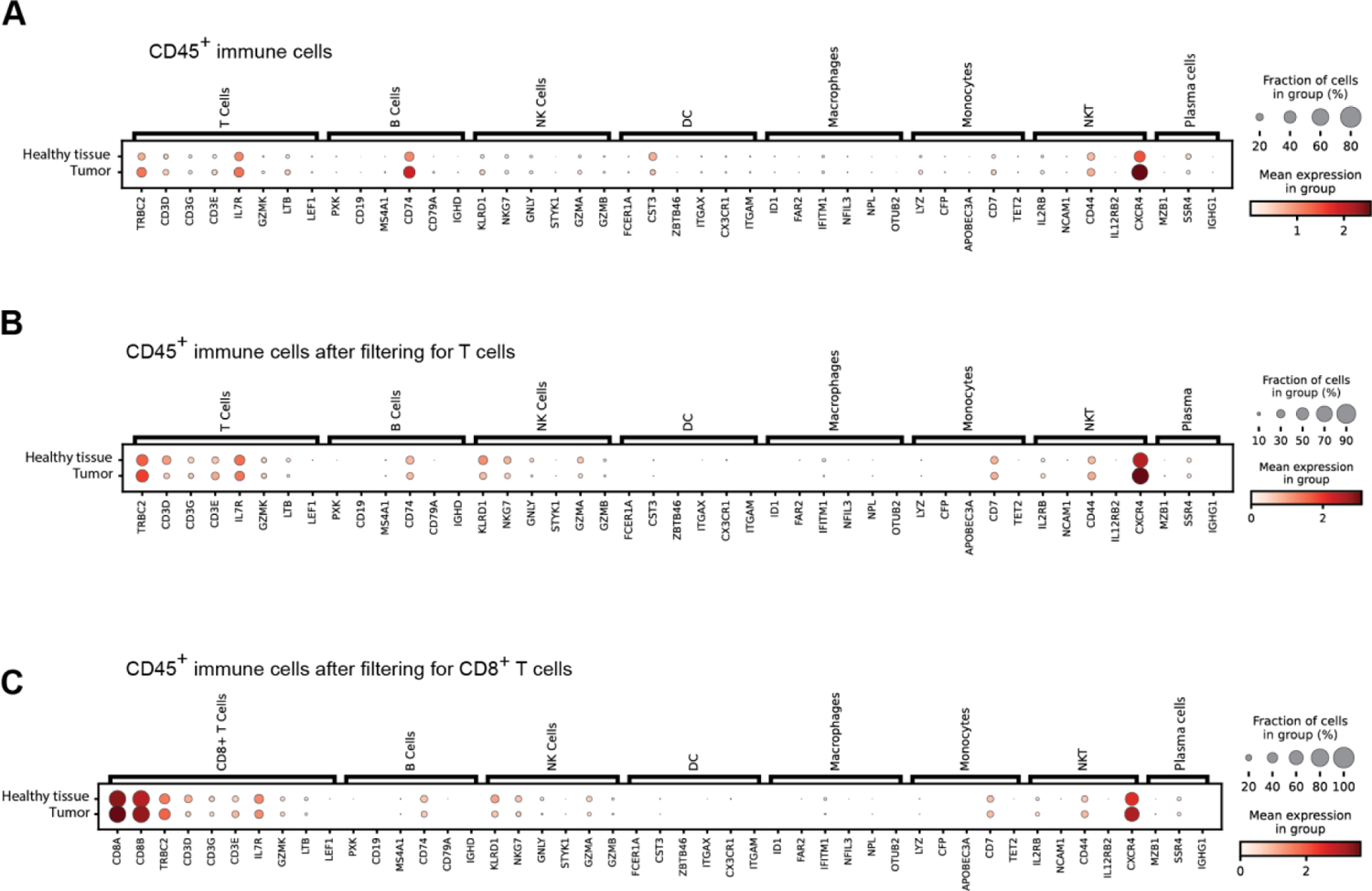
Single-cell transcriptomic annotation of CD8^+^ T cells from breast tumor and healthy peritumoral tissues. **(A)** Marker gene expression for all CD45^+^ immune cells from the matched tumoral and peritumoral tissues from three patients with breast cancer. **(B**, **C**) Marker gene expression for T cells (B) and CD8^+^ T cells (C) from the matched tumoral and peritumoral tissues from three patients with breast cancer after filtering of these immune cell subsets using Celltypist.

**Figure S3.**
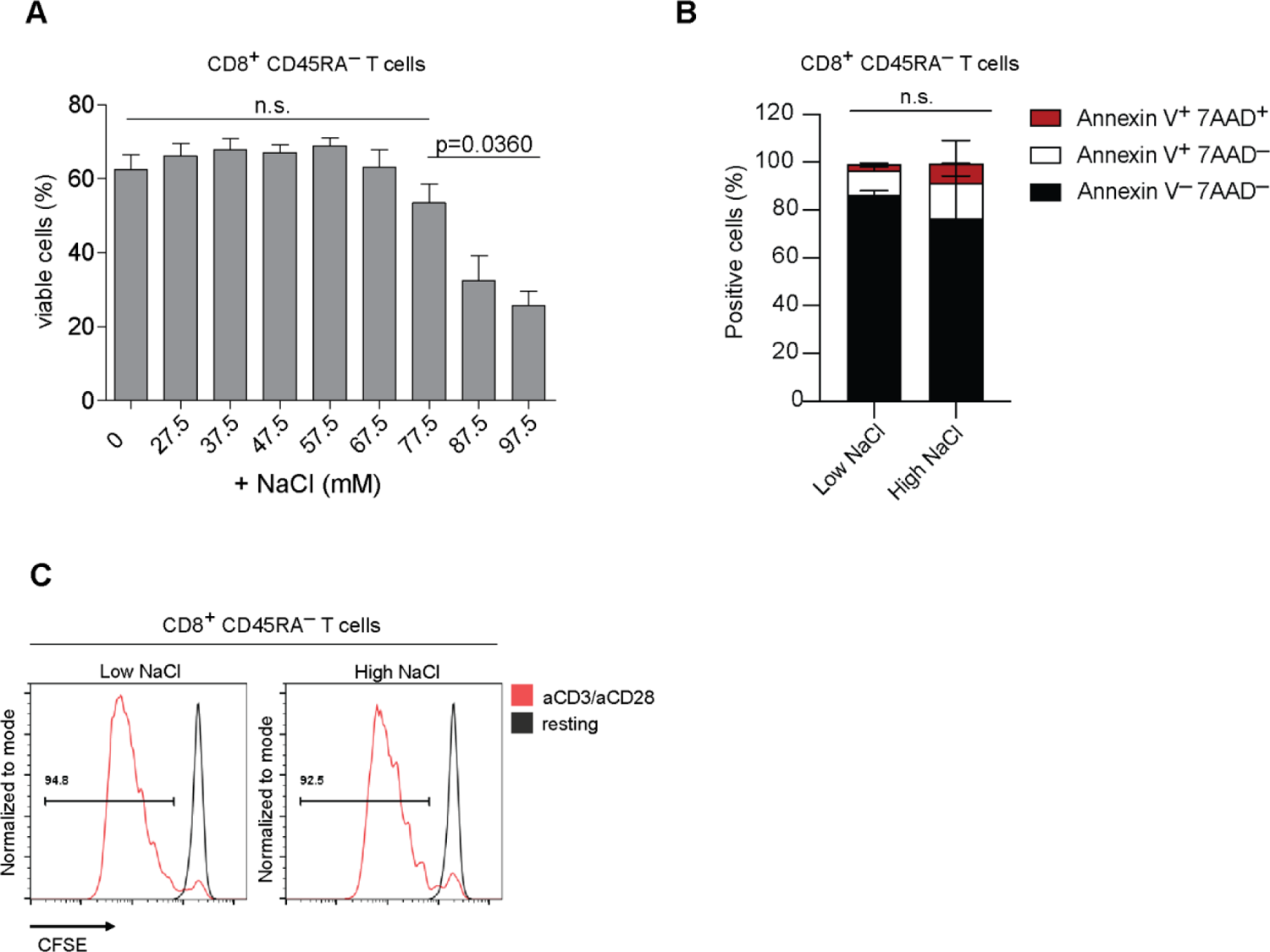
The viability of human CD8^+^ memory T cells is maintained over a wide range of extracellular NaCl concentrations. **(A)** Human CD8^+^ CD45RA^-^ T cells were stimulated with CD3 and CD28 mAbs for 5 days in the presence of increasing (titrated) extracellular NaCl concentrations. Flow cytometric analysis of viability according to live/dead marker dye exclusion. n=3 blood donors. Mean ± SEM. One-way ANOVA. **(B)** Flow cytometry analysis of the indicated markers on human CD8^+^ CD45RA^-^ T cells stimulated as in (A). n=3. One-way ANOVA. **(C)** Flow cytometric analysis of proliferation by CFSE dilution in cells stimulated as in (A). Representative experiment (n=3).

**Figure S4.**
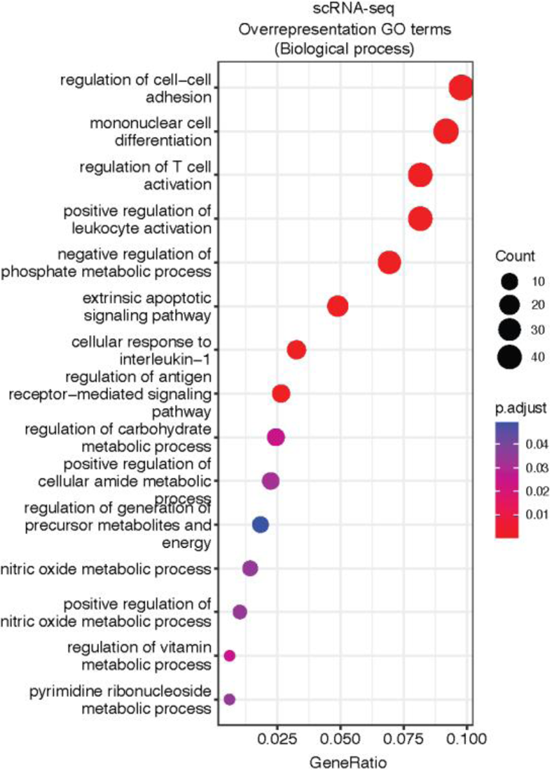
CD8^+^ memory T cells from high NaCl as compared to low NaCl conditions were enriched for transcriptomic processes, indicating invigorated immune responses. scRNA-seq analysis of human CD8^+^ memory T cells stimulated for 3 days under high versus low NaCl conditions was performed. Unbiased overrepresentation analysis using ClusterProfiler. The top 15 processes within the GO terms for “Biological Process” are shown.

**Figure S5.**
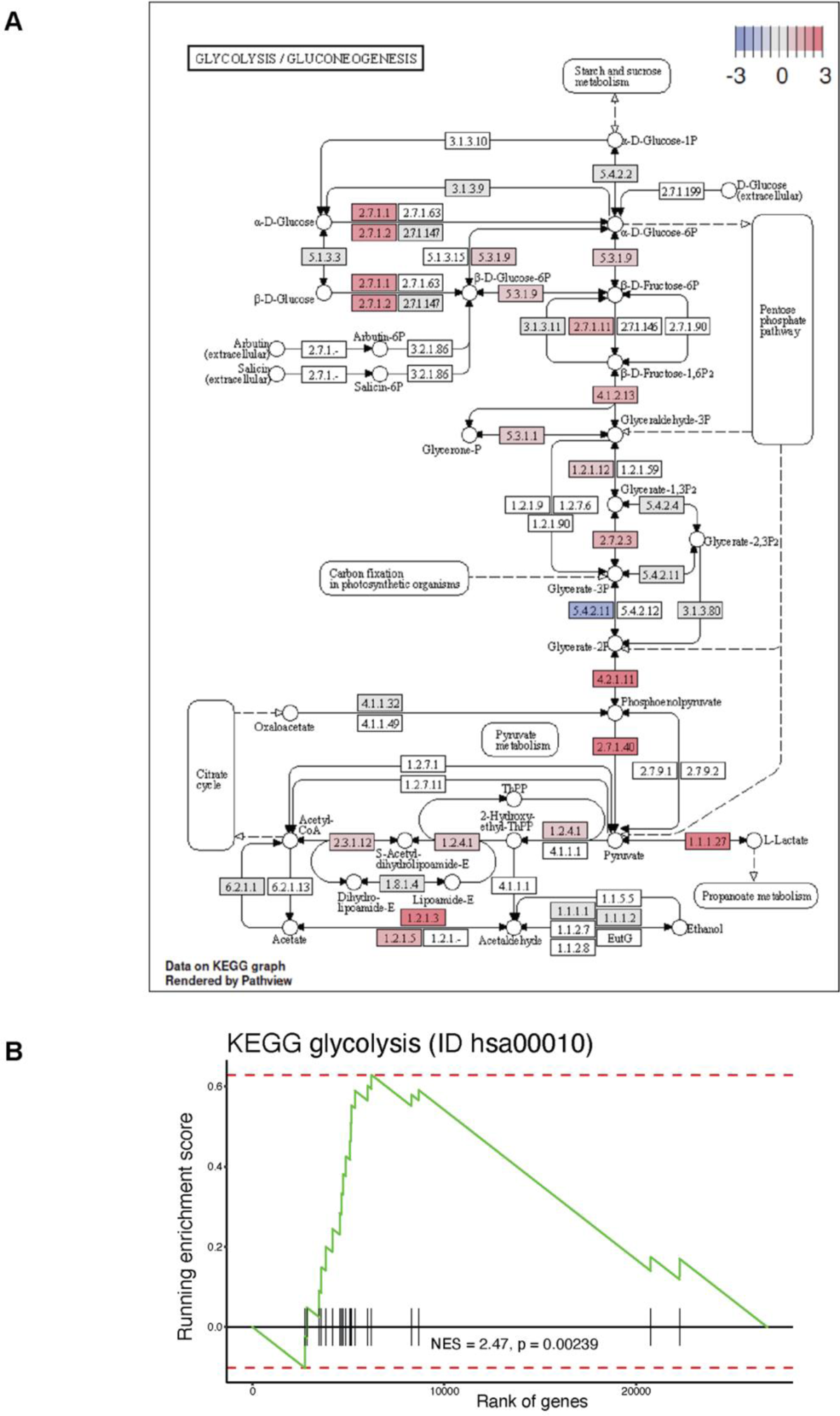
High NaCl conditions promote glycolysis in human CD8^+^ memory T cells on the transcriptomic level. **(A)** KEGG pathway analysis. Expression levels of the individual glycolysis-associated genes following bulk mRNA-seq of human CD8^+^ memory T cells from high versus low NaCl conditions are shown. DESeq2, p≤0.05 **(B)** GSEA of glycolysis associated genes (KEGG pathway: glycolysis). Significance by permutation test (p=0.00239)

**Figure S6.**
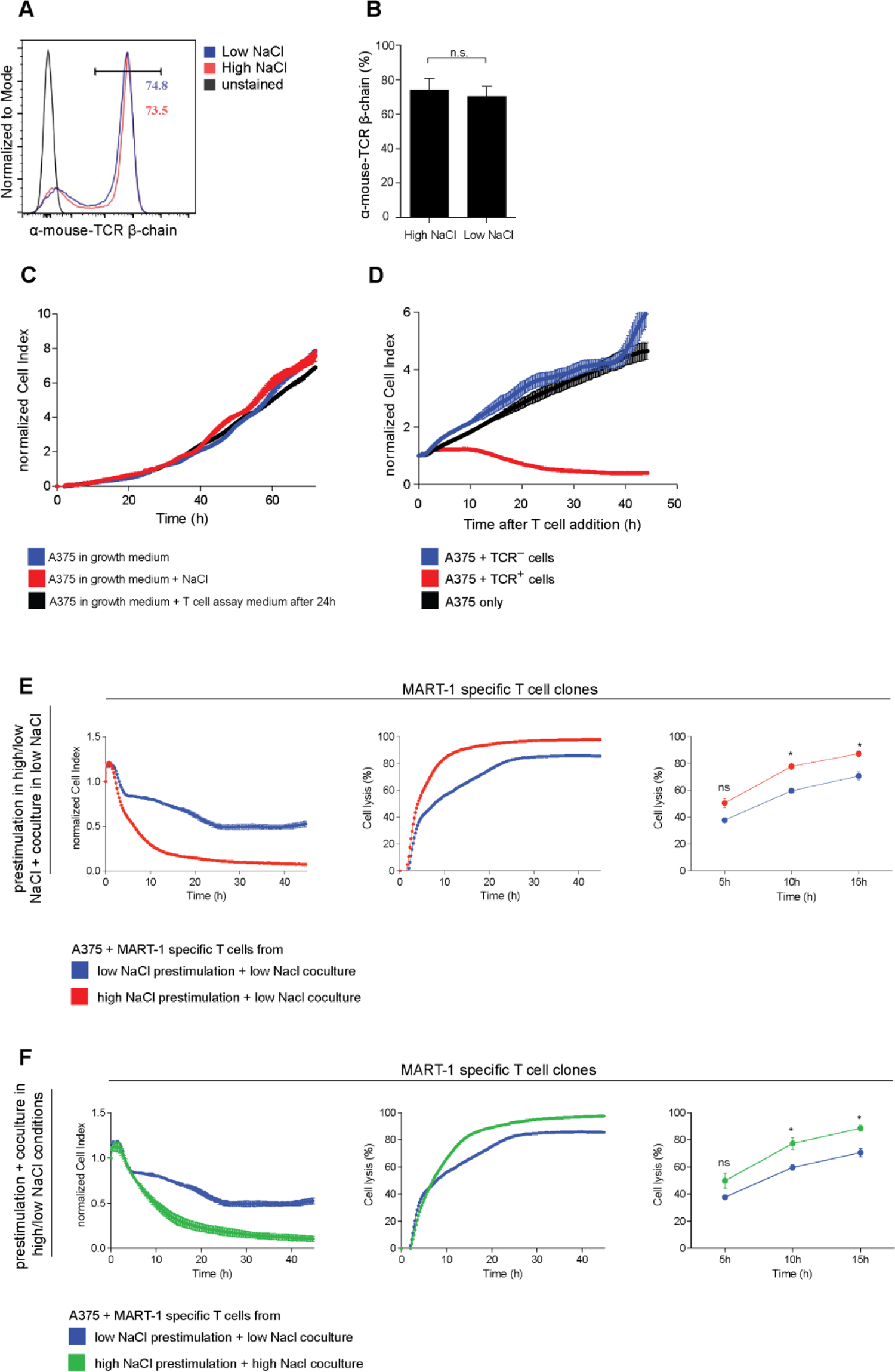
NaCl-induced enhanced killing of target cells by CD8^+^ T cells is antigen specific and not confounded by direct effects of NaCl on target cells. **(A, B)** Flow cytometry of sorted CD8^+^ T cells, which were expanded after nucleofection with a MART-1-specific TCR for 5 days with CD3 and CD28 mAbs under high versus low NaCl conditions. (A), Representative experiment. B, Cumulative quantification (mean ± S.E.M., n=3). **(C)** A375 melanoma cell line cells were seeded on E-plates, and growth was monitored for 72 h under the indicated conditions. One representative experiment is shown (n=4). **(D)** Coculture of MART-1-specific (TCR^+^) or MART-1-nonspecific CD8^+^ T cells (TCR^-^, α-mouse TCRβ chain negative after nucleofection) with A375 melanoma cells (1:1 ratio) pulsed with MART-1 peptide under high versus low NaCl conditions. One representative experiment is shown (n=4). **(E, F)** Coculture of MART-1-specific CD8^+^ T-cell clones with A375 melanoma cells (1:1 ratio) pulsed with MART-1 peptide in the presence or absence of additional NaCl (E) or only absence of NaCl (F) after prestimulation for 5 days with CD3 and CD28 mAbs under high versus low NaCl conditions (n=2 experiments, mean ± S.E.M).

**Figure S7.**
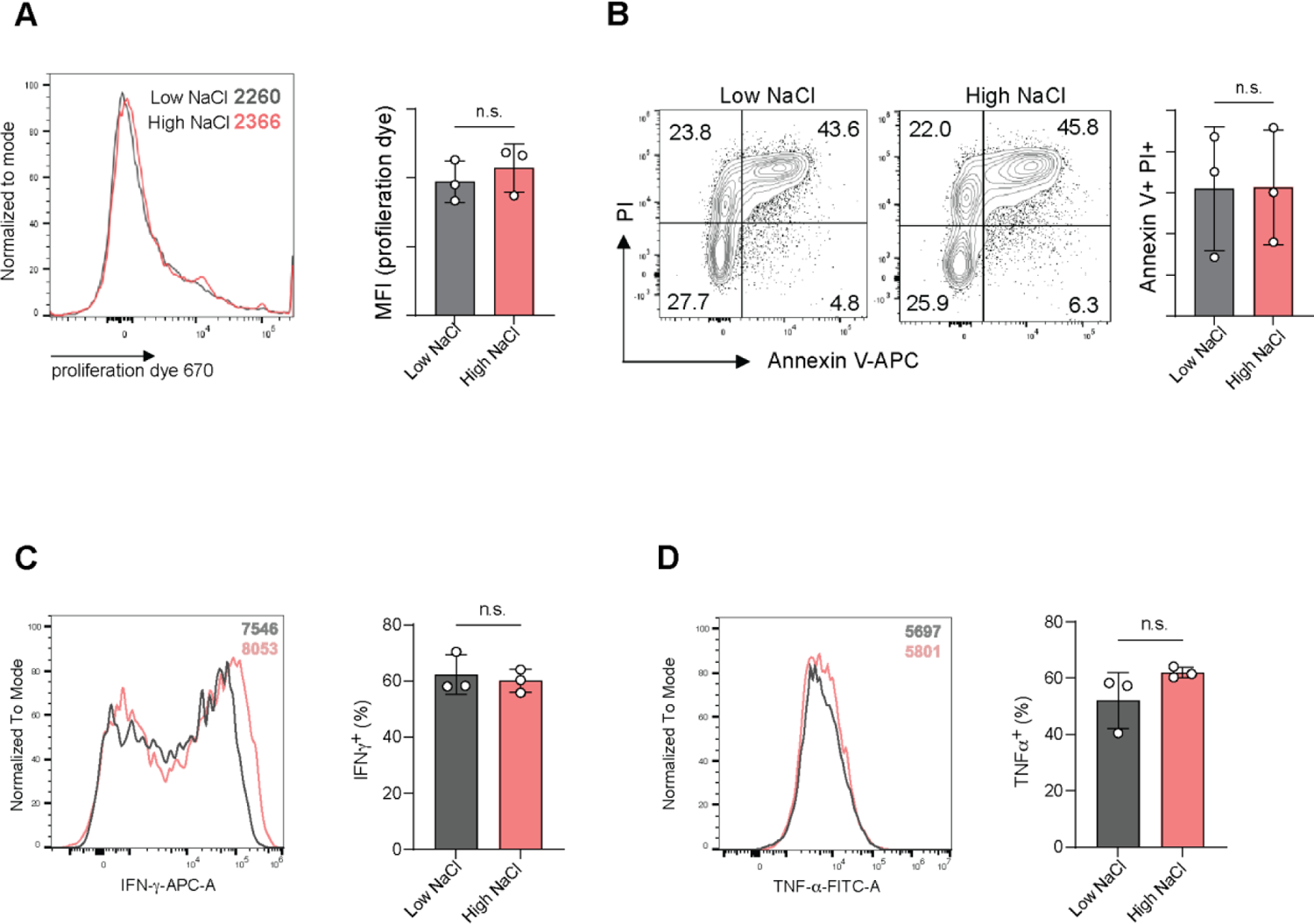
High NaCl conditions do not compromise the viability of mouse CD8^+^ T cells. **(A, B)** Flow cytometric analysis of CD8^+^ T cells after stimulation with CD3 and CD28 mAbs for 3 days under high versus low NaCl conditions (n=3). Left, representative experiment with mean fluorescence intensities shown; right, cumulative quantification. Paired Student’s t test. **(C, D)** Intracellular staining and flow cytometry for IFN-γ (C) or TNF-α (D) on day 3 after stimulation with CD3 and CD28 mAbs for 3 days and restimulation with PMA/ionomycin in the presence of brefeldin A. Paired Student’s t test (n=3). All data points represent biological replicates. n.s., not significant

**Figure S8.**
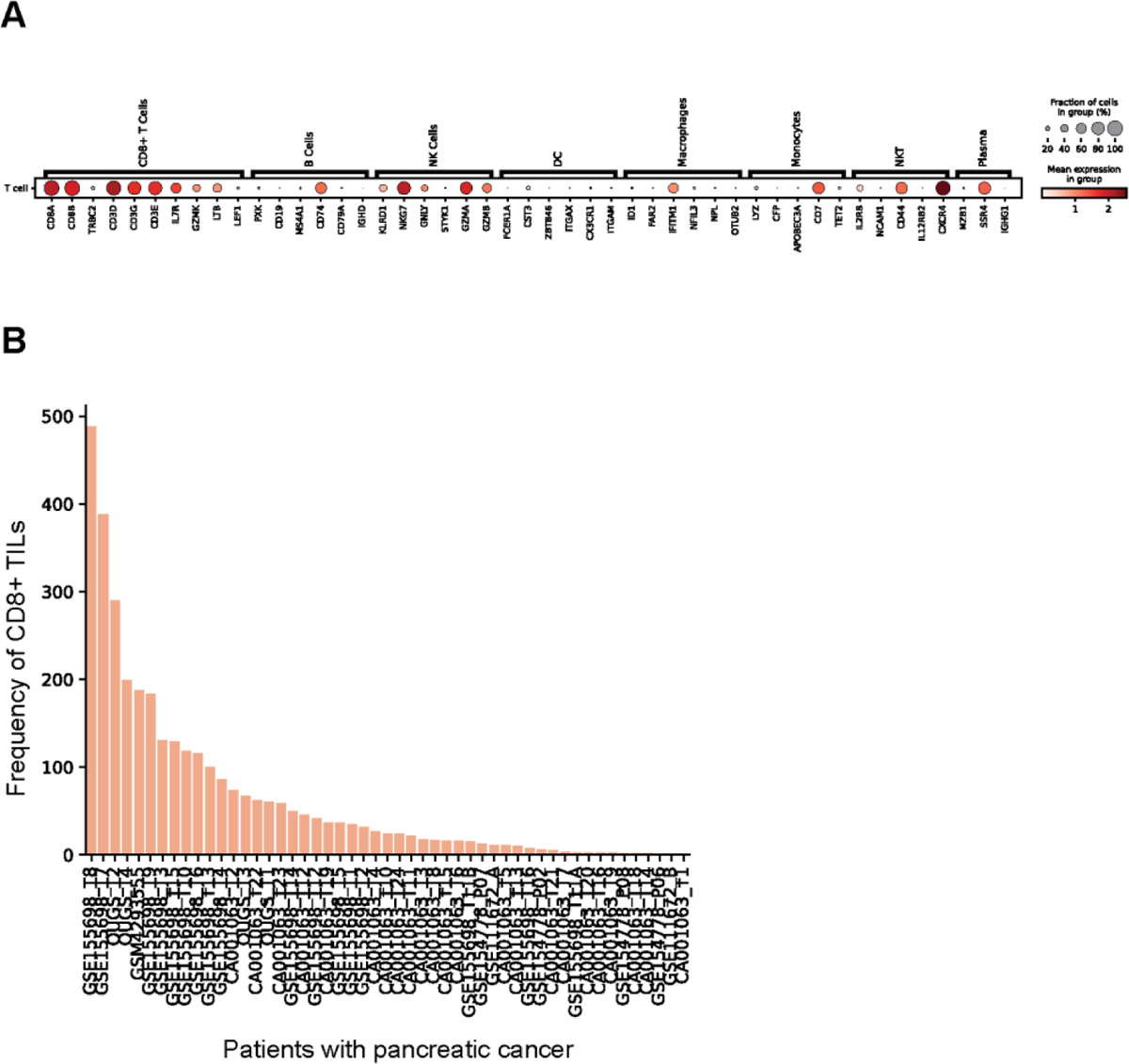
Identification of intratumoral CD8^+^ T cells in whole tumors from patients with pancreatic cancer with scRNA-seq. CD8^+^ T cells from 51 patients with pancreatic cancer identified in scRNA-seq data sets (integration of all cells: 10.5281/zenodo.6024273 (Chijimatsu et al., 2022)). **(A)** Shown is the mean expression of different immune cell lineage marker genes after filtering for CD8^+^ T cells from the whole tumor single-cell transcriptomes. **(B)** Shown is the frequency of CD8^+^ T cells per patient.

## Acknowledgements

We thank all members of the Zielinski lab for fruitful discussions and technical support, particularly Julie Petry, Julia Matthias, Hanna Meinl, Amy Burrell and Ivan Shevchenko. The results shown in Fig. 1D, Fig. 6F and Figure S1 are based in part on data generated by the TCGA Research Network: https://www.cancer.gov/tcga)

## Author Contributions

Conceptualization: C.E.Z. Methodology (bioinformatic): I.G-R., M.A., M.A.A., S.Sc. Methodology (experimental): D.S., C-F.C., S.Su., T.N., P.S., B.M, V.L. Supervision: C.E.Z., M.H., G.P., W.S., H.S. Funding acquisition: C.E.Z., W.S. Writing – review and editing: C.E.Z, D.S., M.A., M.A.A., S.Sc., B.M., V.L., I.G-R., T.N. and P.S., G.P, W.S., M.H., H.S.

